# A rapidly evolving actin mediates fertility and developmental tradeoffs in *Drosophila*

**DOI:** 10.1101/2020.09.28.317503

**Authors:** Courtney M. Schroeder, Sarah A. Tomlin, John R. Valenzuela, Harmit S. Malik

## Abstract

Most actin-related proteins (Arps) are highly conserved in eukaryotes, where they carry out well-defined cellular functions. *Drosophila* and mammals also encode divergent non-canonical Arps in their male-germline whose roles remain unknown. Here, we show that Arp53D, a rapidly-evolving *Drosophila* Arp, localizes to fusomes and actin cones, two male germline-specific actin structures critical for sperm maturation, via its non-canonical N-terminal tail. Although we expected that *Arp53D* loss would reduce male fertility, we instead find that *Arp53D*-KO males are more fertile, both in isolation and in competition with wildtype males. Upon investigating why evolution would retain a gene that negatively impacts male fertility, we unexpectedly found that *Arp53D-KO* females are less fertile. Moreover, KO embryos exhibit reduced viability, which worsens under heat stress. We conclude that ‘testis-specific’ *Arp53D* is detrimental to male fertility, but is required for female fertility and early development, leading to its long-term retention and recurrent adaptation in *Drosophila*.

## Introduction

Actin is an ancient, highly conserved protein that performs many cytoplasmic and nuclear functions vital for eukaryotes, including division, motility, cargo transport, DNA repair and gene expression^1-3^. Its origin predates eukaryotes^4,5^; both bacteria and archaea encode actin-like proteins^6,7^. Actin forms many protein-protein interactions, including with other actin monomers, to perform its various functions^1^. Because of its interactions and functional importance, actin evolves under stringent evolutionary constraints^4,5^. For example, despite being separated by 800 million years of evolution, *D. melanogaster* and *Homo sapiens* actin proteins are 98% identical. In addition to actin, most eukaryotes encode an expanded repertoire of actin-related proteins (Arps) due to ancient gene duplications^4,5^. The resulting Arps have specialized for a wide range of functions, including regulation of actin (Arps 2/3)^8^, chromatin remodeling (Arps 4-8)^9-11^, and microtubule-based transport (Arps 1 and 10)^12,13^. Although different Arps maintain a conserved actin fold, they have specialized for their novel roles via distinct structural insertions^14,15^. These ‘canonical’ Arps significantly diverged from each other early in eukaryote evolution, but now evolve under stringent evolutionary constraints, similar to actin.

Many eukaryotic genomes also encode evolutionarily young, rapidly evolving ‘non-canonical’ Arps. Unlike actin and canonical Arps, which are ubiquitously expressed, non-canonical Arps appear to be exclusively expressed in the male germline^16^. The first described ‘non-canonical’ Arp was *D. melanogaster Arp53D* (named for its cytogenetic location), which was shown to be most highly expressed in the testis^17^. Its presence only in *D. melanogaster* and its unusual expression pattern led to *Arp53D* being mostly ignored in studies of cytoskeletal proteins even in *Drosophila*. However, phylogenomic surveys reveal that ‘non-canonical’ Arps are not as rare as previously believed. Recently, we described a 14-million-year-old *Drosophila* clade that independently acquired four non-canonical *Arp* genes, which are all expressed solely in the male germline^18^. Mammals also encode at least seven non-canonical Arps predominantly expressed in the testis^19-23^, at least some of which localize to actin structures in sperm development^21,22^. This accumulating evidence suggests that non-canonical Arps play fundamentally distinct cytoskeletal functions from canonical Arps, which might explain both their tissue-specificity as well as their unusual evolution.

To gain insight into non-canonical Arps’ functions, we performed evolutionary, genetic, and cytological analyses of *Arp53D* in *D. melanogaster*. We showed that *Arp53D* is strictly conserved over 65 million years of *Drosophila* evolution, suggesting it performs a critical function. Unlike actin or canonical Arps, we found that *Arp53D* has evolved under positive selection. Our cytological analyses reveal that Arp53D specifically localizes to the fusome and actin cones, two specialized actin structures found in the male germline. Arp53D’s unique 40 amino acid N-terminal extension (relative to actin) is necessary and sufficient to recruit it to germline actin structures. Its abundant expression in testes, together with its specialized localization, led us to hypothesize that *Arp53D* loss would lower male fertility. Contrary to this prediction, we found that *Arp53D* knockouts (KO) exhibit increased male fertility. *Arp53D*’s detrimental effect on male fertility is at odds with long-term retention in *Drosophila*. Seeking to explain this paradox, we investigated whether *Arp53D* also had functions outside the male germline. Despite its low expression in females and early embryos, we find that loss of *Arp53D* leads to decreased female fertility and lower embryonic viability under heat stress. Population cage experiments confirm that *Arp53D* has a net fitness benefit in populations despite conferring a fertility loss onto males. We hypothesize that different fitness optima for *Arp53D* for male fertility versus female and embryonic fitness may have resulted in its higher rate of evolution across *Drosophila*. Our study finds that non-canonical ‘testis-expressed’ Arps can be highly conserved over broad evolutionary spans and play critical roles outside the male germline.

## Results

### *Arp53D* encodes a rapidly evolving non-canonical Arp that has been strictly retained for over 50 million years

*Arp53D* was first identified as a male-specific Arp gene in *Drosophila melanogaster*^17^. It was subsequently shown to be phylogenetically more closely related to *actin* rather than to any of the canonical Arps^4^, suggesting it arose via a more recent gene duplication from *actin. Arp53D* was not found in any other non-insect genomes in a broad survey of eukaryotes, raising the possibility that it only exists in a few *Drosophila* species. To date its evolutionary origin, we investigated *Arp53D* presence in sequenced genomes from *Drosophila* and closely related *Diptera*^24-28^. We found that *Arp53D* has been strictly retained for over 50 million years of *Drosophila* evolution^24^ (Figure 1A, Table S1). Deleterious or non-functional genes are quickly pseudogenized and lost within a few million years in *Drosophila* genomes^29^. Thus, its strict retention implies that *Arp53D* has been selectively retained for an important function in *Drosophila* despite its more recent evolutionary origin.

**Figure 1.**
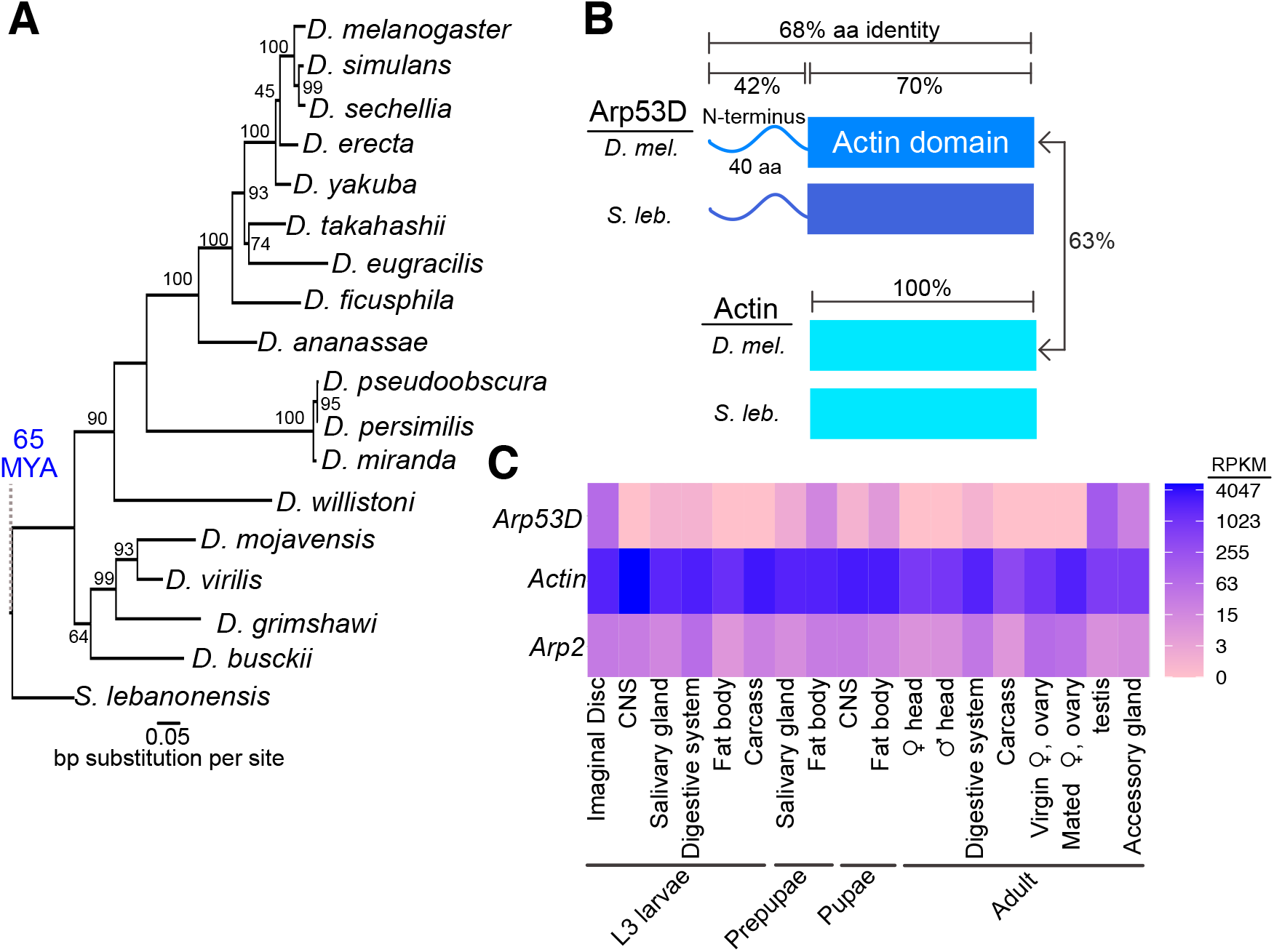
*Arp53D* encodes a rapidly evolving non-canonical Arp with male-enriched expression. **A)** Nucleotide tree with 100 bootstraps includes *Arp53D* orthologs from sequenced genomes of 18 *Drosophila* species. **B)** Arp53D domains include an extended N-terminus, which is predicted to be unstructured, and the canonical actin domain. The protein identities are displayed for the different domains of actin (Act5C) and Arp53D from the species *D. melanogaster* and *Scaptodrosophila lebanonensis*. **C)** RNA-seq values (in RPKM) are displayed for tissues at different developmental stages (wandering L3 larvae, white prepupae, pupae, and adults)^36,70^. Values are displayed for *Arp53D, Act5C*, and *Arp2* with blue indicating highest expression.

To gain insight into its function, we compared the domain architectures of *D. melanogaster* Arp53D to Act5C, one of two non-muscle actins encoded in *Drosophila* genomes. Like canonical Arps, Arp53D includes an actin fold domain, which consists of 4 subdomains and a central ATP-binding pocket (Figure 1B). What distinguishes Arp53D from actin is an extended 40 amino acid N-terminal domain that is predicted to be mostly unstructured (Figure 1B, Supplementary Figure S1A). All *Arp53D* orthologs encode this extended N-terminal domain, which is also the most rapidly evolving segment of Arp53D in sequence and length. For example, N-terminal domains from *D. melanogaster* and *S. lebanonensis* Arp53D proteins are only 42% identical whereas the actin fold domain is 70% identical (Figure 1B). In contrast to Arp53D, actin homologs are 100% identical over a comparable period of evolutionary divergence.

Since actin evolves under extremely strong selective constraint, the higher divergence of *Arp53D* could simply reflect more relaxed selective constraints. Alternatively, it could reflect a faster than expected divergence of *Arp53D* due to diversifying selection. To distinguish between these possibilities, we took advantage of publicly available sequences of hundreds of *D. melanogaster* strains^30,31^ (www.popfly.org^32^) to carry out McDonald-Kreitman (MK) tests for positive selection^33^. The MK test compares the ratio of fixed non-synonymous (amino acid replacing, P_N_) to synonymous (P_s_) polymorphisms within a species (*D. melanogaster*) to fixed differences between species (D_N_ and D_S_, *D. melanogaster-D. simulans)*; we exclude low frequency polymorphisms since they have not been subject to selective scrutiny^34,35^. If selective constraints are not significantly altered within versus between species, we expect D_N_:D_S_ to be approximately equal to P_N_:P_s_. We analyzed several actin and Arp genes using the MK test. Unfortunately, several actin genes have no non-synonymous changes (fixed or polymorphic) and cannot be suitably analyzed. The MK test found no evidence of positive selection in several canonical Arp homologs (Supplementary Figure S1B). In contrast, we found an excess of D_N_:D_S_ (21:24) compared to P_N_:P_s_ (1:19) for *Arp53D*, implying it has evolved under strong diversifying (positive) selection during the *D. melanogaster-D. simulans* divergence (p=0.001) (Supplementary Figure S1B). The non-synonymous fixed differences between species are distributed throughout the entirety of the gene with most changes concentrated toward the N-terminal half (Supplementary Figure S1C). Our evolutionary analyses thus find that *Arp53D* is subject to atypical selective constraints for an Arp in *D. melanogaster*, consistent with it performing a distinct function from canonical Arps.

### Arp53D localizes to specific actin structures late in sperm development

*Arp53D* was first shown to be expressed in *D. melanogaster* testes^17^. We took advantage of transcriptomic profiling of various adult tissues in *D. melanogaster* and nine other *Drosophila* species to investigate which tissues express *Arp53D*. Confirming previous analyses, we found that all *Drosophila* species show significantly male-biased expression of *Arp53D* and undetectable expression in adult females (Supplementary Figure S1D, Table S2). In all these cases, *Arp53D* RNA expression is much higher in the testis than the remaining male carcass (Supplementary Figure S1D, Table S2). More detailed and extensive transcriptome profiling of various tissues in *D. melanogaster*^36^ revealed that *Arp53D* RNA expression is also modestly expressed in other tissues, including fat bodies and imaginal discs at earlier developmental stages (Figure 1C). This extremely sex-and tissue-biased expression of *Arp53D* is highly unusual for an *actin* or canonical *Arp* gene, which are ubiquitously expressed in all tissues (Figure 1C).

We investigated Arp53D localization in *D. melanogaster* testes, where it is most abundantly expressed. *Drosophila* testes contain numerous cell types, including somatic cells and germ cells at many stages of development (*i*.*e*., mitotic cells, meiotic cells, and mature sperm)^37^. Germ cells undergo incomplete cytokinesis during their four mitotic divisions and subsequent meiosis, resulting in a cyst of 64 sperm cells, which share the same cytoplasm and membrane until full maturation^37^ (Figure 2A). Multiple cysts at different stages of spermatogenesis are visible in the testis at any given time, allowing simultaneous visualization of all developmental stages.

**Figure 2.**
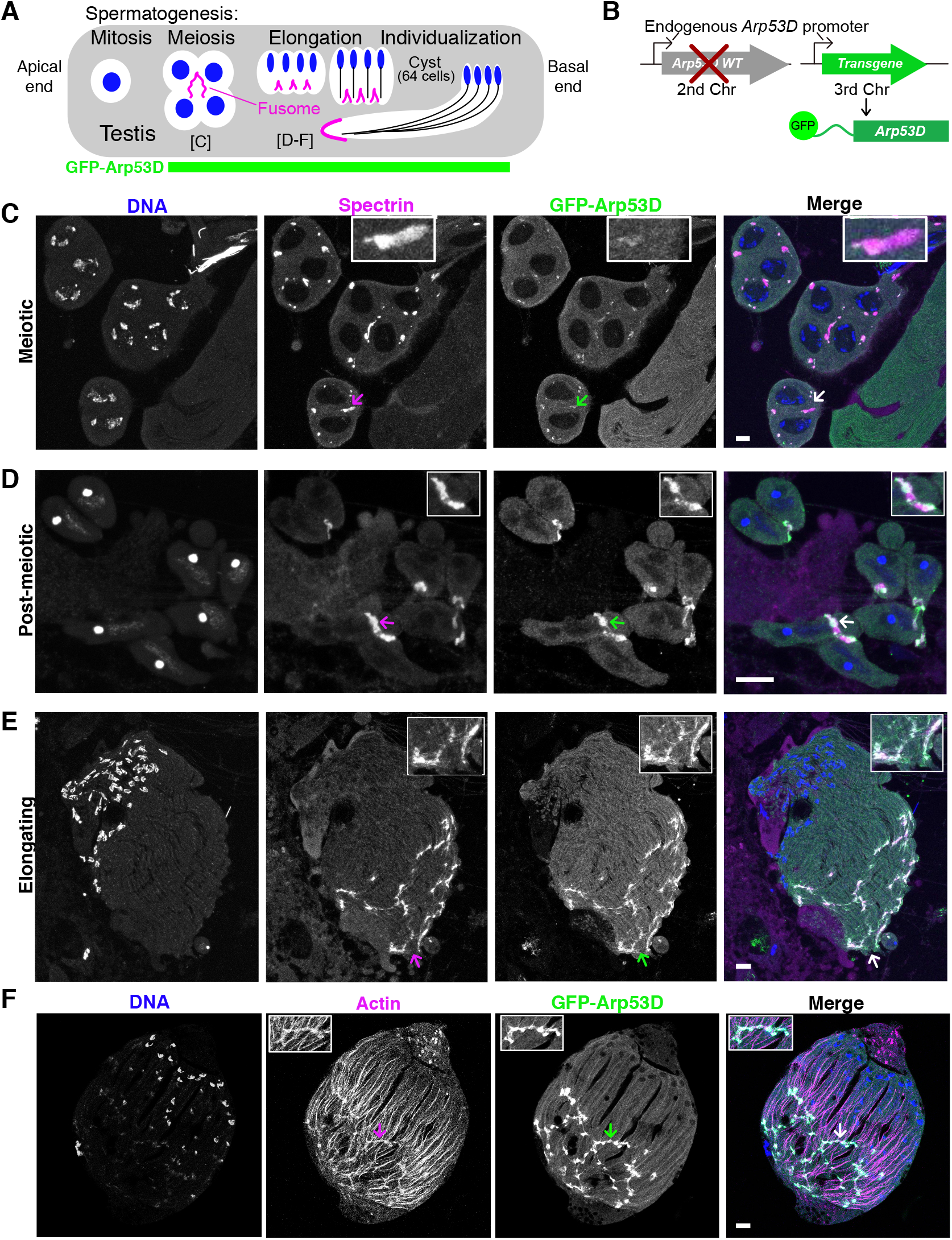
Arp53D localizes to specific actin structures late in sperm development. **A)** A schematic shows the steps of spermatogenesis that progress from the apical end to the basal end of the testis; green indicates the level of GFP fluorescence visualized in the testis of the transgenic fly in (B). Labels for the meiotic and elongating stages refer to panels C-F. **B)** A transgene encoding Arp53D with an N-terminal superfolder GFP (sfGFP) tag was site directed to the third chromosome. The transgenic fly line was then crossed into the Arp53D-KO background so that all Arp53D molecules are fluorescent. **C-E)** Testes from the transgenic flies in (B) were broken apart and cysts from meiotic (C), post-meiotic (D) and elongating stages (E) of spermatogenesis were visualized and probed for the fusome-localizing protein α-spectrin (magenta), DNA (blue), and Arp53D (green, anti-GFP). The merge of spectrin and Arp53D appears as white. Arrows correspond to the enlarged insets. **F)** Testes from the transgenic flies in (B) were fixed and probed for actin (phalloidin). All scale bars are 10 µm.

We generated a fly transgenic line with superfolder GFP-tagged *Arp53D* (*sfGFP-Arp53D*) under the control of its endogenous promoter (Figure 2B). This transgene was introduced and assayed in an *Arp53D*-knockout background such that only two copies of *sfGFP-Arp53D* are present. Thus, every Arp53D molecule is fluorescently tagged, allowing for improved detection. We found that sfGFP-Arp53D is undetectable during mitosis but is present within the meiotic and post-meiotic spermatocyte cysts (Figure 2, Supplementary Figure S2A-B) where it localizes specifically to two male germline-specific actin structures: the fusome during meiosis and spermatid elongation (Figure 2) and actin cones during sperm individualization (Figure 3).

**Figure 3.**
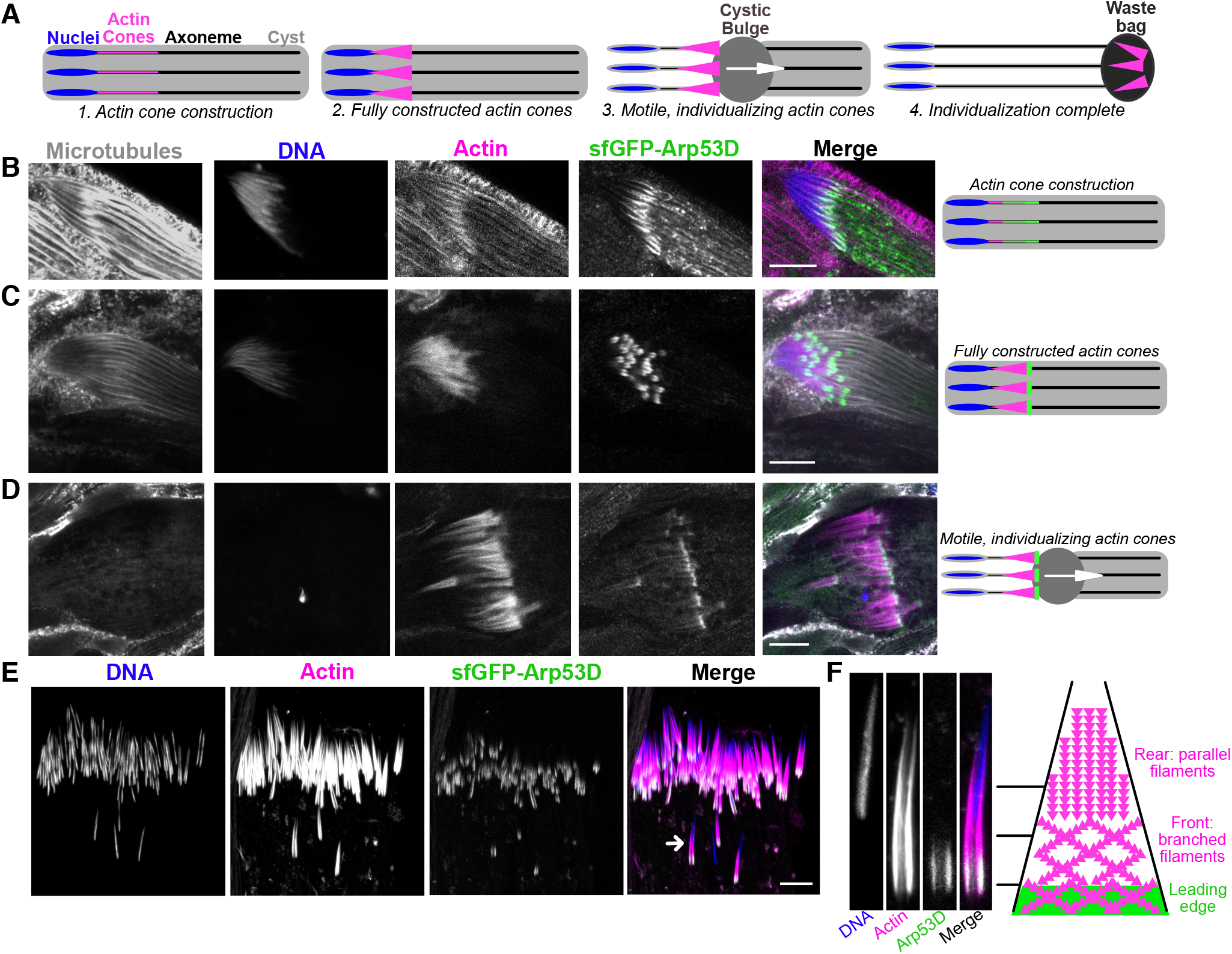
Arp53D localizes to the leading edge of actin cones during sperm individualization. **A)** The schematic depicts the stages of individualization. Once actin cones are fully assembled at mature sperm nuclei, the cones translocate along the axoneme (a microtubule structure), pushing excess cytoplasm (the “cystic bulge”) to the end of the cyst. The cystic bulge undergoes autophagy and becomes known as the “waste bag.” **B-D)** Testes from the same flies in Figure 2B were dissected and fixed. Microtubules (gray, anti-tubulin), DNA (blue, DAPI), actin (magenta, phalloidin), and sfGFP-Arp53D (green, anti-GFP) were visualized. Each row shows a cyst at a different stage of individualization, which is depicted with a schematic to the right. **E)** Testes expressing sfGFP-Arp53D (Figure 2B) were dissected and imaged live. Actin (sir-actin probe) and mature sperm nuclei (Hoechst stain) were visualized in addition to GFP. The arrow indicates the actin cone shown in (F). **F)** A mature sperm nucleus and its corresponding actin cone is shown with Arp53D localizing at the leading edge. On the right is a schematic that delineates the types of actin networks composing the cone. The green filaments indicate Arp53D localization. All scale bars are 10 µm.

The fusome is an actin-coated membranous organelle that forms at all incomplete cytokinetic furrows following mitosis and meiosis. It forms a large network that connects all developing spermatids, mediating cytoplasm exchange within the cyst^38,39^ (Figure 2A, Supplementary Figure S2C). To ascertain sfGFP-Arp53D localization to the fusome, we fixed sfGFP-Arp53D-expressing testes and probed for the fusome-specific α-spectrin protein^40^. We found that a majority of sfGFP-Arp53D co-localizes with α-spectrin, confirming Arp53D localization to the fusome (Figure 2C-E). Arp53D localization to the fusome is weak during meiosis (Figure 2C) but becomes progressively stronger post-meiosis, at which stage sfGFP-Arp53D localizes at increasing intensity to the entirety of the fusome (Figure 2D-E). Arp53D remains associated with the fusome even as it moves to one end of an elongating cyst. We conclude that Arp53D is targeted to the actin-coated fusome specifically during meiosis with increased recruitment to the fusome during spermatid elongation. Arp53D’s localization specifically to the fusome is in contrast to that of actin localization, which occurs throughout the cyst (Figure 2F, Supplementary Figure S2C).

During late stages of spermatogenesis, spermatids must separate and obtain their own individual membranes. This process is carried out by the ‘actin cone’, a cone of actin filaments, which forms around each sperm head when nuclear condensation is complete. Cones translocate along the axoneme of the sperm tail to push out excess cytoplasm (“cystic bulge”) while encasing each sperm in its own membrane^41,42^ (Figure 3A). All 64 actin cones move synchronously down the spermatid tails and then undergo degradation along with the excess cytoplasmic components in what becomes known as the “waste bag” ^41,42^ (Figure 3A). When actin cones begin to polymerize (indicated by a gradual accumulation of filamentous actin), we find that sfGFP-Arp53D is linearly enriched along the axoneme and slightly overlaps the base of sperm nuclei (Figure 3B). At this stage, sfGFP-Arp53D localization is indistinguishable from actin. However, when actin cones are fully formed, sfGFP-Arp53D is visible as a highly concentrated punctate structure at the front of the actin cone, distinct from actin (Figure 3C, E). Subsequently, sfGFP-Arp53D remains associated with actin cones as they translocate even when they pass the axoneme near the end of the cyst (Figure 3D). The leading edge of the actin cone where sfGFP-Arp53D localizes is composed of branched actin networks (Figure 3E-F) and is the site of active actin polymerization and membrane extrusion, whereas the rear of the actin cone is composed of parallel actin bundles^43^ (Figure 3F). Previous studies have shown that an actin-binding molecular motor—myosin VI—also localizes to the leading edge of actin cones^44^. Indeed, we find that a testis-specific myosin VI subunit^45^ colocalizes with Arp53D at the leading edge (Supplementary Figure S2D). Proteomic studies^46^ and our cytological analyses (Supplementary Figure S2E) do not detect Arp53D in mature sperm. We, therefore, conclude that Arp53D protein must be degraded in the waste bag with the rest of the actin cone apparatus.

Thus, Arp53D specifically localizes to two germline-specific actin structures in a dynamic manner. It first localizes to the fusome as it forms during meiosis (Figure 2) and then moves to actin cones as they are being constructed once spermatid elongation is complete (Figure 3). It remains associated with actin cones until it is ultimately destroyed along with the rest of the actin cones following the completion of sperm individualization. Notably, for most of spermatogenesis, Arp53D localization is distinct from actin, which localizes more broadly than Arp53D. Our cytological analyses suggest that Arp53D carries out specialized roles at unique cytoskeletal machineries during male reproduction.

### Arp53d’s unique N-terminal extension is necessary and sufficient for recruitment to germline cytoskeletal structures

We investigated whether Arp53D’s unique 40-residue N-terminal domain mediated its specialized localization to the fusome and actin cones (Figure 1B). We generated a *sfGFP-△N-term D. melanogaster* transgenic line encoding *sfGFP-Arp53D* with 35 amino acids of the N-terminal domain deleted (Figure 4A). The *sfGFP-△N-term* transgene was driven by the endogenous *Arp53D* promoter from the same insertion site in the fly genome as our full length *sfGFP-Arp53D* transgene (Figure 2B). We dissected testes from the transgenic flies and performed immunoblotting analyses, which showed that the smaller deletion protein was being expressed at comparable levels as sfGFP-Arp53D (Supplementary Figure S3B). Moreover, similar to full-length *sfGFP-Arp53D* transgenic flies, we could show that *sfGFP-△N-term* transgenic flies also express GFP in meiosis (Supplementary Figure S3A). However, unlike full-length Arp53D, the GFP localization remained diffuse, and we did not detect concentrated GFP signal at the fusome or actin cones (Figure 4B-C, Supplementary Figure S3C). Based on these experiments, we conclude that the N-terminus is necessary for Arp53D’s localization to these specialized germline actin structures. We can further conclude that Arp53D’s actin fold domain is divergent enough that it cannot co-polymerize with actin *in vivo*.

**Figure 4.**
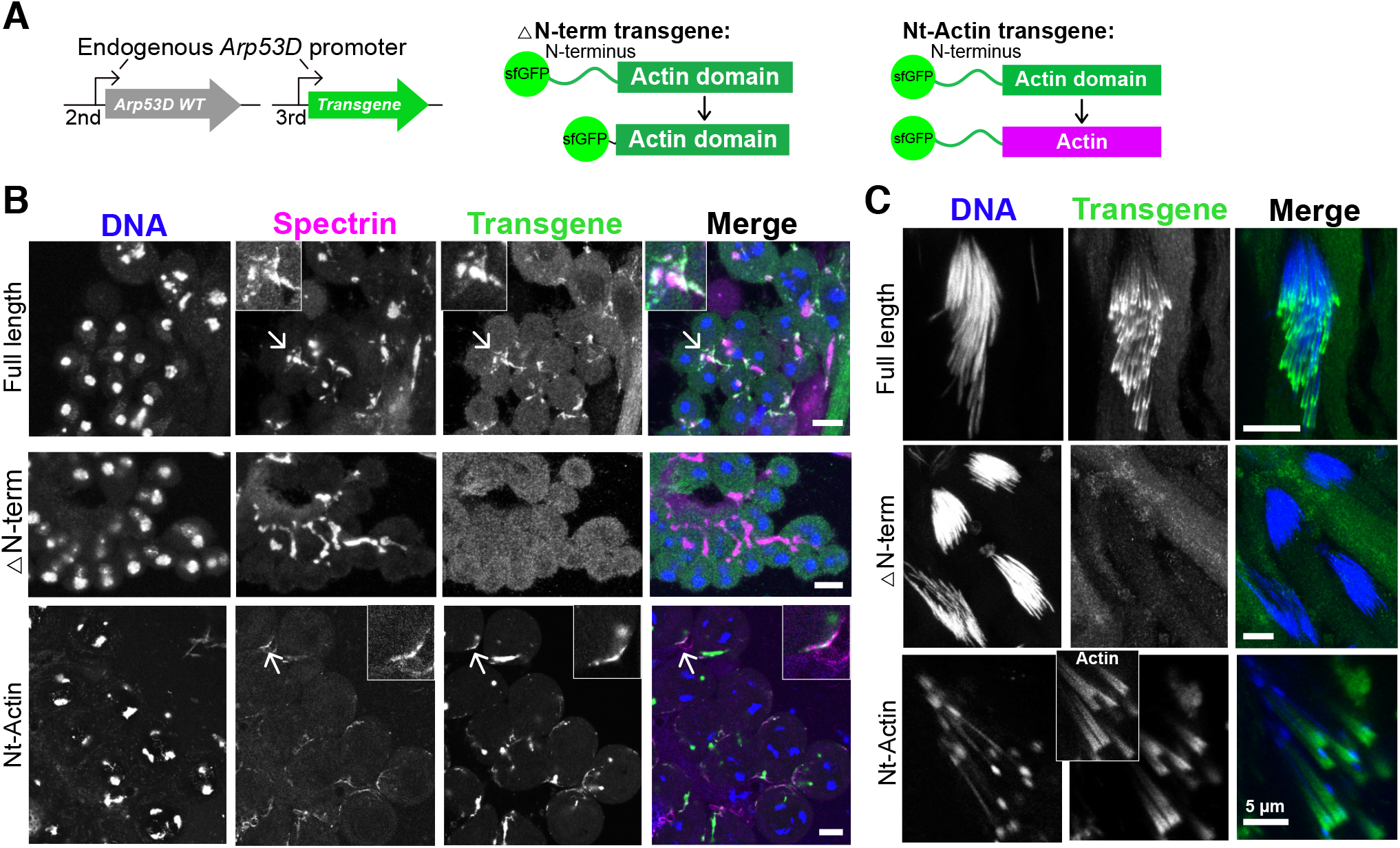
N-terminus of Arp53D is necessary and sufficient for localization. **A)** Two transgenic fly lines were generated with the transgene on the third chromosome in the WT background. In the “△N-term” transgene, 35 aa of the N-terminus of Arp53D was removed and the remaining actin fold was N-terminally tagged with sfGFP. In the “Nt-Actin” transgene, the actin domain of Arp53D was replaced with canonical actin (Act5C). Both transgenes were under the control of *Arp53D’s* endogenous promoter. **B)** Dissected testes of the transgenic flies were fixed and probed with anti-GFP (green), anti-spectrin (magenta) and Hoechst (blue). Arrows correspond to the enlarged insets. All scale bars are 10 µm. **C)** Dissected testes of the transgenic flies were fixed and probed to amplify signal with anti-GFP (green) and Hoechst (blue), and mature sperm nuclei were imaged. Nt-actin was imaged live, and the inset shows labeled actin (sir-actin probe). All scale bars are 10 µm unless otherwise noted.

We next tested whether the N-terminus was sufficient to confer Arp53D’s localization to canonical actin. We generated an *sfGFP-Nt-actin D. melanogaster* transgenic line, encoding sfGFP-Arp53D N-terminal domain-canonical actin (Act5C) (Figure 4A). Like all previous transgenic constructs, we placed this chimeric protein under the control of *Arp53D*’s endogenous promoter and kept the same genomic insertion location (Figure 4A). We found that this chimeric protein is expressed and localizes similarly to full-length sfGFP-Arp53D throughout spermatogenesis, maintaining its association with the fusome during spermatid elongation and motile actin cones throughout individualization (Supplementary Figure S3D) just like full-length Arp53D (Figure 4B-C). Furthermore, despite encoding an identical actin fold domain, this chimeric protein did not appear to co-colocalize with actin throughout the developing cysts. Based on these findings, we conclude that the most prominent structural diversification of Arp53D—its N-terminal extension—is both necessary and sufficient for recruitment of actin to unique male cytoskeletal machinery.

However, the N-terminal domain cannot confer this specialized localization onto any globular protein. When we tested the localization of Arp53D’s N-terminal domain with solely sfGFP (“N-term-sfGFP”, Supplementary Figure S3E), we could only detect diffuse GFP expression and no concentrated signal at the fusome or actin cones (Supplementary Figure S3E). This implies that specialized localization to fusomes and actin cones requires both the Arp53D N-terminal domain as well as the actin fold domain from actin or an actin-like protein.

### Loss of *Arp53D* increases male fertility

Based on its strict retention in *Drosophila* and its cytological localization to germline-specific actin structures in *D. melanogaster* testes, we predicted that *Arp53D* must play important roles in male fertility. To test this hypothesis, we created a knockout (KO) of *Arp53D* using CRISPR/Cas9, introducing an early stop codon and a *DsRed* transgene under the control of an eye-specific promoter (Supplementary Figure S4A). The *DsRed* transgene allowed us to track the KO allele by fluorescence microscopy and to distinguish heterozygous from homozygous KO flies based on intensity of eye fluorescence. We backcrossed the KO founder line to a wild type strain (Oregon-R) for 8 generations in order to isogenize the KO background with Oregon-R as much as possible (Supplementary Figure S4B). We sequence-verified the presence of *DsRed* in the *Arp53D* locus (Supplementary Figure S4C-E) and lack of mutations in the upstream essential gene *SOD2*. We also confirmed lack of *Arp53D* expression in the KOs and absence of *Wolbachia*, a bacteria that can infect wildtype strains of *Drosophila* and confound fertility assays^47^ (Supplementary Figure S4F-G).

One possible consequence of Arp53D loss in KO males is that we might expect to see gross disruption of the germline actin structures to which it localizes. Contrary to this expectation, we found no gross defects in overall organization or actin intensity of actin cones or the fusome in *Arp53D-KO* males (Supplementary Figure S4H-I). We reasoned that loss of Arp53D may have led to more subtle organizational defects that would manifest in fertility reduction of *Arp53-KO* males. To evaluate any fertility defects, we mated WT females to either homozygous *Arp53-KO* males or isogenic WT males for 9 days and subsequently counted all progeny that survived to adulthood (all crosses are written female x male, Figure 5A). We were surprised to find that the KO males had significantly higher fertility compared to WT males at 25°C (1.3-fold increase in average progeny count, Figure 5B, p=0.001) or an even more pronounced advantage at 29°C (2.4-fold difference in the average progeny count, Figure 5B, p<0.0001). We also found that this reduction in fertility is dose-dependent; heterozygous *Arp53D*-KO males have intermediate fertility between homozygous KO and WT males (Supplementary Figure S5A). Thus, presence of only one intact copy of *Arp53D* is sufficient to reduce male fertility at 29°C (p=0.03), while two copies are significantly worse (p<0.0001).

**Figure 5.**
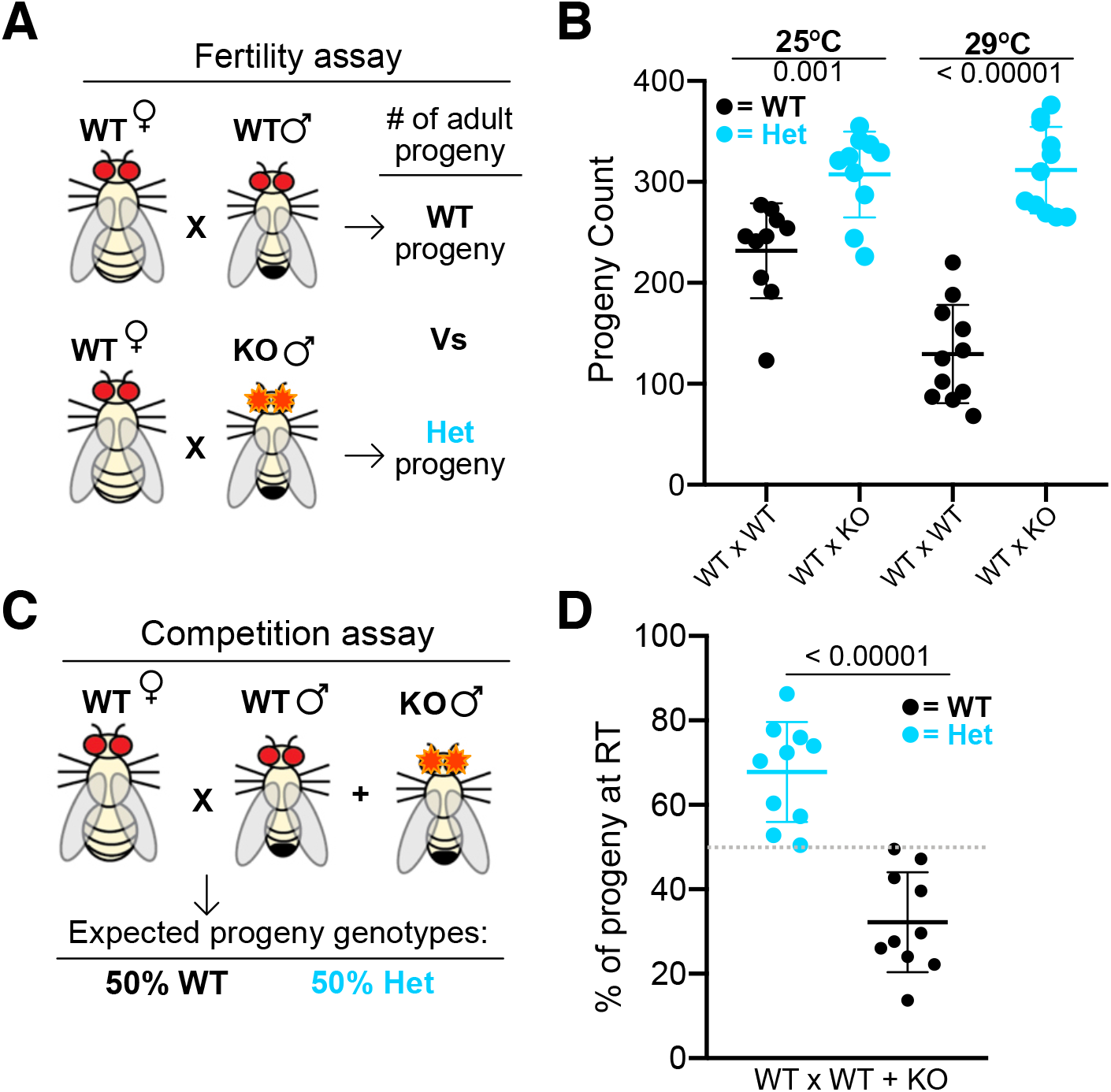
Lack of Arp53D increases male fertility. **A-B)** Fertility assays at 25°C and 29°C were conducted with WT females mated to either WT males or KO males (all crosses are reported as female x male). Embryos were laid for 9 days and adult progeny were counted. **C-D)** WT females were mated to WT males and KO males at 25°C. Progeny of KO and WT males were identified with presence or lack of DsRed fluorescence, respectively. Progeny genotypes are displayed as a percentage of the total population. For all graphs, progeny genotypes are distinguished by color, and a one-way ANOVA test was used to determine all p-values.

We were concerned about the possibility that the CRISPR/Cas9 approach may have resulted in an off-target linked mutation that had escaped several rounds of isogenization. We therefore adopted an orthogonal approach to validate our surprising findings about the increased fertility of *Arp53D*-KO males. We conducted RNAi knockdown of *Arp53D* using topi-Gal4^48^ to induce expression of the RNAi hairpin specifically targeted against the *Arp53D* coding region (Supplementary Figure S5B-C). Consistent with our genetic KO, we found that knockdown of *Arp53D* resulted in significantly increased fertility at 29°C (p=0.005, Supplementary Figure S5C). Together with the *Arp53D*-KO results, these data strongly argue that lack of *Arp53D* increases male fertility.

Our assays of male fertility so far were all done in the absence of competition between males. We hypothesized that although *Arp53D* intrinsically decreases male fertility, it may confer a competitive advantage in the presence of other males, which might explain its long-term retention in *Drosophila*. To test this possibility, we mated WT females to both WT males and *Arp53D*-*KO* males (Figure 5C). If WT and *Arp53D*-*KO* males had equal probabilities of successfully fertilizing, then 50% of adult progeny would be fathered by WT or *Arp53D-KO* males (Figure 5C). However, we found that *Arp53D*-*KO* males sired nearly 70% of the progeny even in the presence of WT males, implying that they had a significant fertility advantage even in a competitive situation (p<0.0001, Figure 5D). Our experiments show that *Arp53D* had a significant, although unexpectedly deleterious, effect on male fertility both in isolation and in competitive scenarios, leaving unanswered the question of why it was strictly retained in *Drosophila*.

### Loss of *Arp53D* has deleterious consequences to female fertility and embryonic viability

Given that *Arp53D* presence was disadvantageous to male fertility, we considered whether other life history traits require *Arp53D*, which might help explain its long-term evolutionary retention. Although *Arp53D* is most abundantly expressed in adult testes, there is weaker expression in other tissues and developmental stages (Figure 1C). Although we could not detect any *Arp53D* expression in adult females (Figure 1C), we still wanted to rule out an effect of *Arp53D* on female fertility. To investigate this possibility, we crossed *Arp53D*-KO females to either WT males or *Arp53D*-KO males, and compared the number of adult progeny produced relative to WTxWT crosses (Figure 6A). At room temperature (25°C), we did not observe any significant differences between any of these three crosses (Figure 6B). Since KO males are more fertile than WT males (Figure 5), we might have expected to see increased number of adult progeny in the KOxKO crosses, but we did not (Figure 6B). This suggested the possibility that there was some fitness impairment in the KOxKO crosses, that could have arisen either because of the maternal or zygotic genotype.

**Figure 6.**
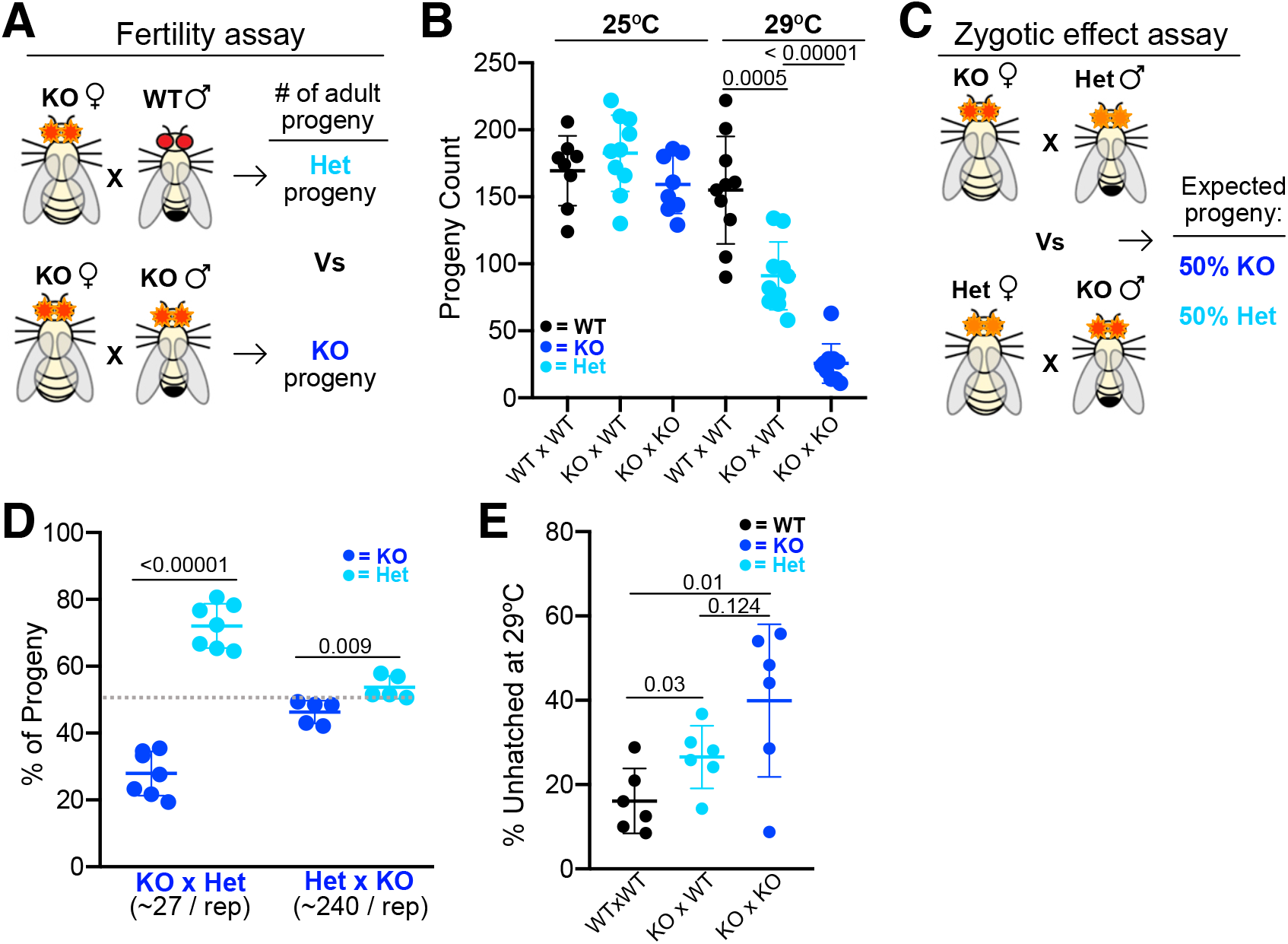
Lack of Arp53D in females leads to developmental tradeoffs. **A-B)** Fertility assays at 25°C and 29°C were conducted with KO females mated to either WT males or KO males. Embryos were laid for 9 days and adult progeny were counted. **C-D)** Arp53D-KO females were crossed to Het males, and Het females were crossed to KO males. Homozygous KO progeny were identified with strikingly bright DsRed fluorescence, whereas the Het progeny were identified by dim fluorescence. Progeny genotypes were quantified and displayed as a percentage of the total population. **E)** Crosses were allowed to lay for 2 hr at 29°C, and embryos were counted. Unhatched embryos were then quantified 24 hours later and are displayed as a percentage of embryos laid the previous day. A one-way ANOVA test was used to determine all p-values.

At 29°C, this impairment became much more obvious. We found that the number of adult progeny that are produced in KOxWT crosses, which produce heterozygous (HET) progeny, was significantly reduced relative to WTxWT crosses, which produce WT progeny (p=0.0005, Figure 6B). Based on these results, we infer that *Arp53D-KO* females have a clear fertility disadvantage, which is exacerbated at higher temperatures. The number of adult progeny produced in KOxKO crosses was even further reduced 3.5 fold relative to KOxWT crosses (p<0.0001, Figure 6B), in spite of the fact that KO males are more fertile than WT males (Figure 5B). We attribute this dramatic reduction in progeny survival to the zygotic genotype, since only KO progeny can be produced from KOxKO crosses.

To further distinguish the effects of maternal versus zygotic loss of Arp53D, we conducted two additional crosses—HETxKO and KOxHET—at 29°C, to accentuate any fitness differences (Figure 6C). In both crosses, the progeny produced are either KO or HET. If there were no contribution of zygotic genotype on fitness, we would predict that both crosses should result in approximately 50% KO progeny (dotted line, Figure 6D). Instead, we found that KO progeny were recovered at slightly lower than 50% frequency in the HETxKO crosses (p-value<0.01, Figure 6D), suggesting that the KO progeny are at a fitness disadvantage compared to the HET progeny. This difference in fitness was significantly exacerbated in the KOxHET crosses, in which KO progeny only made up <30% of total progeny (p<0.0001, Figure 6D). Thus, both crosses show that KO zygotes are at a fitness disadvantage relative to HET zygotes. However, the even stronger effect in the KOxHET crosses relative to the HETxKO crosses on both fraction of KO progeny produced (Figure 6D) and total progeny produced (Supplementary Figure S5D) further implicates maternal genotype as playing a more significant deleterious effect on progeny survival than paternal genotype.

We wished to further understand the nature of the defects associated with loss of fitness upon zygotic loss of *Arp53D*. We therefore conducted three crosses in parallel at 29°C: WTxWT, KOxWT, and KOxKO. We saw no significant differences in the number of eggs laid between these sets of crosses (Supplementary Figure S5E). However, in the KOxKO crosses, we found that a large percentage of eggs failed to develop (Figure 6E). Whereas <20% of eggs failed to develop to larval stages in WTxWT crosses, nearly 40% of eggs failed to develop in the KOxKO crosses (p=0.01, Figure 6E). The bulk of this difference can be explained due to defects in female fertility or maternal contribution, which results in significant differences between the WTxWT and KOxWT crosses (p=0.03, Figure 6E). In contrast, we found more modest differences between KOxKO and KOxWT crosses (p=0.124, Figure 6E), which would result from differences in zygotic genotype. Overall, we conclude that both maternal and zygotic *Arp53D* genotypes contribute to optimal embryonic fitness in *D. melanogaster*. Thus, unexpectedly, we find that not only does *Arp53D* have a negative effect on male fertility (Figure 5), but it is also required in females and zygotes for optimal fitness (Figure 6).

The significant genetic contribution of maternal and zygotic *Arp53D* to fitness is perplexing because it had very low to undetectable levels of expression in previous transcriptomic studies^36^. However, bulk RNA-seq analyses can miss transcripts that are expressed at low levels. To address this discrepancy, we carried out additional studies to examine *Arp53D* expression. First, we carried out RT-PCR analyses with a greater degree of sensitivity (more amplification cycles), which revealed that Arp53D is indeed expressed in adult females albeit at much lower levels than males (Supplementary Figure S6A). Although previous RNA-seq data does show very low levels of Arp53D expression in the ovary^49,50^, our cytological examination of ovaries in female flies expressing *sfGFP-Arp53D* did not reveal GFP expression above background levels (Supplementary Figure S6B-C). We also investigated embryonic expression of Arp53D by examining existing *in situ* data for *D. melanogaster* gene expression. This analysis revealed weak expression of *Arp53D* in stages 1-3 of embryogenesis (Supplementary Figure S6D)^51^, suggesting maternal contribution. Furthermore, single cell-transcriptome data of embryos show localized *Arp53D* RNA in stage 6 embryos (Supplementary Figure S6E)^52^. Interestingly, Arp53D localizes dorsally in stage 6 embryos whereas canonical actin is concentrated ventrally (*Act5C*, Supplementary Figure S6E), further demonstrating functional divergence of *Arp53D* from its parental gene. Single-embryo RNA-seq also indicates zygotic expression of *Arp53D*^53^. Thus, overall, despite weak expression, we conclude that Arp53D is sufficiently expressed in embryos to manifest its zygotic effects, whereas the maternal tissues in which *Arp53D* is expressed is still unclear.

### *Arp53D* retention provides an overall fitness advantage in *D. melanogaster*

Our analysis of fitness effects of *Arp53D* reveal two opposing phenotypes. Loss of *Arp53D* is beneficial to male fertility, but detrimental to female fertility and zygotic viability. Our finding that Arp53D is strictly retained for over 50 million years of *Drosophila* evolution suggests that its positive consequences must outweigh its negative consequences. To explicitly test this possibility, we competed KO and WT alleles of *Arp53D* over multiple generations using a population cage experiment. In this assay, we used Arp53D-KO flies that were isogenized in a *w1118* genetic background (6 backcrosses) due to the lack of eye pigmentation and, thus, more efficient identification of DsRed fluorescence. These *D. melanogaster* strains are completely isogenic except for the loss of *Arp53D* and gain of eye-expressed *DsRed* in the KO allele.

We began the experiment with 3 starting populations consisting of an excess of KO flies (75%) to put the *Arp53D* KOs at a starting advantage (Figure 7A). At each generation, we randomly selected 50 females and 50 males (without observing the fluorescent eye marker for the *Arp53D-KO* allele) and quantified the remaining progeny for the presence of the *Arp53D-KO* allele (Figure 7B). After 20 generations at room temperature, we found a robust and consistent increase of the WT allele, which finally reached an average proportion of 67% in all three replicate populations (Figure 7B). This rise from 25% to 67% frequency in just 20 generations implies a roughly 2% fitness advantage for the WT allele per generation. This implies that there is a significant selective coefficient associated with the retention of *Arp53D* at the population level. Thus, even though males lacking *Arp53D* appear to be at an advantage, this benefit is outweighed by fitness costs attributed to roles in female fertility and embryonic development. Based on these results, we conclude that non-canonical *Arp53D* mediates fitness tradeoffs in order to play important roles in multiple life history traits of *D. melanogaster*.

**Figure 7.**
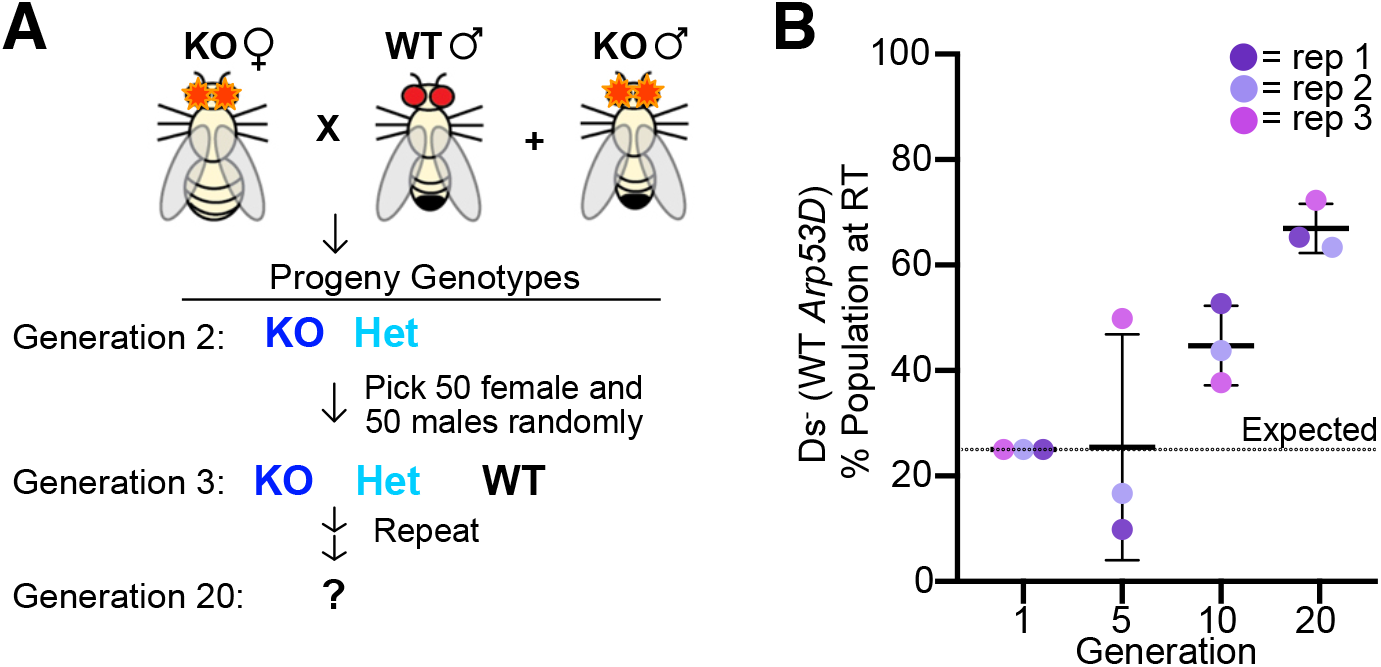
Arp53D exhibits an evolutionary fitness advantage. **A)** A population cage experiment was conducted with generation one including *Arp53D-KO* females, *Arp53D-KO* males, and WT (*w1118*) males in the same bottle. *Arp53D-KOs* were isogenized in the *w1118* background. All subsequent generations were passaged by randomly selecting 100 progeny from the previous generation and placing them in a new bottle at room temperature. **B)** All progeny at generations were assessed for presence of DsRed-fluorescent eyes, the marker for the *Arp53D-KO* allele. The graph displays the percent of WT progeny (flies lacking DsRed fluroescence) of the generation’s total population, and replicates are distinguished by color. The dotted line indicates the expected percentage of WT progeny over time if no fitness advantage is associated with WT Arp53D.

## Discussion

Actin and canonical actin-related proteins (Arps) represent some of the most conserved proteins in eukaryotic genomes. Canonical Arps diversified early in evolution and have been mostly retained for their essential cellular functions since. In contrast to these ancient, conserved Arps, many genomes also encode non-canonical Arps that are often evolutionarily young, rapidly evolving, and predominantly expressed in the male germline. Because these non-canonical Arps have not been retained for large evolutionary periods, show fewer hallmarks of evolutionary constraint, and are predominantly expressed in the transcriptionally promiscuous male germline, they have received less scientific attention than canonical Arps. In this study, we investigated one of the earliest described non-canonical Arps, encoded by *Arp53D* in *D. melanogaster*. Although this Arp is not widely conserved even in animal genomes, we show that it is evolutionarily ancient (at least 65 million years old), strictly retained, and is important for optimal fitness in *D. melanogaster*. Moreover, even though *Arp53D* is predominantly expressed in *Drosophila* testes, we find that it exerts its critical function in other developmental stages including early embryogenesis. Non-canonical Arps like *Arp53D* are found in many animal genomes, including mammals. Our analyses suggest that these previously ignored non-canonical Arps might encode many unusual and distinct functions.

The evolutionary invention of Arps allows the deployment of the actin fold to perform new functions without compromising actin’s many essential functions within a cell. The more recent evolutionary divergence of non-canonical Arps provides a better opportunity to dissect how they diverged from actin to acquire and consolidate their varied cellular functions. For example, Arp53D is distinguished from canonical actin by its rapidly evolving actin-fold domain and a longer 40 amino acid residue N-terminal domain. Although N-terminal tails in actin proteins are typically much shorter---only 3 amino acid residues in length---they regulate the binding of many regulatory proteins, such as myosin^54,55^ and are often post-translationally modified to affect actin localization, polymerization and interactions with actin-binding proteins^56^. Our analyses show that the longer N-terminal tail is necessary and sufficient to explain Arp53D’s specialization to germline-specific actin structures during spermatogenesis. We hypothesize that the unique, longer N-terminal tail of Arp53D may similarly allow it the ability to interact specifically with other cytoskeletal proteins, thereby distinguishing it from canonical actin.

It is not unexpected that novel Arps would specialize for spermatogenesis, which requires several novel cytoskeletal functions and complex actin structures. For example, *Drosophila* exhibits a unique sperm developmental program that deploys two germline-specific actin structures: the fusome and actin cones. Arp53D specializes to localize to both in a developmental stage-specific manner that is distinct from actin. Actin cones are unique to *Drosophila* flies, whereas the fusome is found in both female and male flies, additional insects^57^, and also frogs^58^. Although these actin structures are absent in many species’ germ cell developmental programs, the actin-based processes of cytoplasm sharing and sperm separation span many phyla^57,59^. We hypothesize that the requirement of these specialized actin processes may have led to the independent origin and retention of many non-canonical Arps throughout animal evolution. Indeed, we find that another non-canonical Arp that arose via gene duplication from canonical *Arp2* in the *D. pseudoobscura* lineage, independently specialized to localize to actin cones^18^. We speculate that their role in reproduction, as a class, may have led to their accelerated rate of evolution due to strong selective pressures from sperm competition and sexual selection^60-62^.

Given its rapid evolution, predominant expression in male testes and localization to specialized actin structures in spermatogenesis, we expected that loss of *Arp53D* would lead to a significant impairment of male fertility. Instead, we saw the opposite; loss of *Arp53D* led to an increase in male fertility, whether in isolation or in competition with WT males, at 25°C or 29°C. Given this unexpected finding, we wondered why *Arp53D* is even expressed in testes? One possibility is that *Arp53D* may serve to monitor the quality of sperm produced. Thus, despite being fertilization competent, *Arp53D*-KO sperm may lead to progressively less fit progeny. Except for our population cage experiments (Figure 7), our other assays are only one generation long and thus lack the sensitivity to detect any such subtle detrimental effect. Alternatively, other evolutionary pressures that we have not tested in our assay (like the presence of Wolbachia) could potentially reveal a ‘hidden benefit’ for *Arp53D* function in the male germline. Given that a predominant testis-specific expression pattern is a hallmark of *Arp53D* and other non-canonical Arps in *Drosophila* and mammalian species, *Arp53D*’s beneficial role on male fertility remains an open possibility that we cannot completely exclude.

In contrast to the male germline, *Arp53D* clearly plays a beneficial role both in female fertility as well as zygotic viability. Female fertility differences could be on account of mating differences, fertilization competence of the oocyte, or impact on progeny development (maternal effect). We note that the fitness defects arising in KOxWT crosses are significantly less severe than in the KOxKO crosses. This suggests that there is no gross impairment of female fertility in *Arp53D*-KO females. *Arp53D* expression in *D. melanogaster* ovaries cannot be reliably detected either by RNA-seq analyses or by sfGFP-Arp53D visualization (Supplementary Figure S6B-C). However, *in situ* detection of *Arp53D* in early embryos (Supplementary Figure S6D)^51^ suggests that *Arp53D* RNA may be maternally deposited. In contrast to maternal contributions, *in situ* and RNA-seq data show robust *Arp53D* expression in the later stages of development^51,52^, consistent with a zygotic effect. Inviability of *Arp53D*-KO zygotes was significantly rescued if they were from HET females, but not if they were derived from KO females (Figure 6D-E, Supplementary Figure S5D). This raises the possibility that *Arp53D* is a maternal-zygotic lethal effect gene, where small consequences of either maternal effect or zygotic effect lethality are dramatically exacerbated when both maternal and zygotic genotypes are *Arp53D*-KO.

The nature of the zygotic defect caused by loss of Arp53D remains unknown. Given its ability to localize to germline-specific actin structures, Arp53D may be localizing to actin beyond the testis as well. Interestingly, actin networks are drastically reorganized in the heat stress response of *Drosophila* embryos, leading to decreased embryonic viability^63^. We speculate that Arp53D may regulate embryonic actin networks that are important in this process, explaining why a zygotic effect is strongly exacerbated at high temperature.

Our studies reveal that, contrary to assumptions based on patterns of highest expression, non-canonical *Arp53D* plays important roles in many aspects of *D. melanogaster* biology beyond male fertility. Many genes exhibit highest expression in the testis and brain, two tissues that are especially transcriptionally promiscuous, despite the fact that their most important function may manifest elsewhere. The ‘out-of-testis’ hypothesis predicts that the male germline provides an initial ‘gene nursery’ for evolutionary innovation, with diversification subsequently broadening its expression profile^64,65^. A recent cytological study of a novel centromeric histone variant, which was originally thought to be ‘testis-specific’ in *Drosophila virilis*, based on RT-PCR data, demonstrated that its expression and function also extends to females^66^. Similarly, Umbrea which is highly testis-enriched, is required for chromosome segregation more broadly^67^. Thus, although surprising, it is not unprecedented for a testis-enriched gene like *Arp53D* to be retained for its function outside the male germline.

Despite *Arp53D*’s detrimental effect on male fertility, our population cage experiments firmly establish that it confers a net fitness advantage to *D. melanogaster* populations, presumably because of its beneficial consequences on female fertility and zygotic viability. Thus, *Arp53D* appears to be poised in a state where it cannot simultaneously be beneficial to male fertility and female/zygotic fitness. Such a genetic conflict has precedence. For example, while absence of the rapidly evolving transcription factor, Eip74EF, leads to increased male fertility, female fertility is reduced^68^. Under such circumstances, it is expected that this impasse should be resolved by transcriptional rewiring, which can occur rapidly. For example, *Apollo* and *Artemis* nuclear importin genes duplicated and transcriptionally diverged ∼200,000 years ago in the *D. melanogaster* lineage^69^. While *Apollo* is required for male fertility, it is detrimental to female fertility, while the opposite is true for *Artemis*^69^. By analogy, we would predict that males should lose *Arp53D* expression in testes to relieve male fertility defects but maintain female and zygotic expression. Yet, such transcriptional rewiring has not occurred in over 50 million years of *Drosophila* evolution. It is possible that the random mutations required to separate the transcriptional programs are either rare or impossible for *Arp53D*. In such a case, *Arp53D* is trapped to maintain a delicate balance to affect male fertility least deleteriously while enhancing female and zygotic fitness the most. This balance may require recurrent adaptation to optimize. Thus, our studies suggest that a balancing ‘war between the sexes’ may be ultimately driving the rapid evolution of a non-canonical Arp in *Drosophila*.

## Materials and Methods

### Phylogenetics and positive selection tests

All sequences (Table S1) were obtained from Flybase^70^ and/or NCBI and aligned using MAFFT^71^ in Geneious^72^. Protein sequences were used for maximum likelihood trees, generated using PhyML and 100 bootstraps. For positive selection tests, unpolarized McDonald-Kreitman tests^73^ were conducted with 197 *D. melanogaster* strains (DPGP3)^30^ as the ingroup, and the *D. simulans* allele^74^ was used as the outgroup, with rare polymorphisms (<5%) removed^34^. A chi-square test was used to assess statistical significance of the difference between fixed and polymorphic non-synonymous and synonymous changes.

### Sequencing and RT-PCR

To obtain genomic DNA from flies for subsequent PCRs and Sanger sequencing, 1 or 2 flies were ground in 10 mM Tris-HCl pH8, 1 mM EDTA, 25 mM NaCl, and 200 µg/mL Proteinase K. The fly lysate was incubated at 37°C for 30 min, followed by 95°C for 3 min to inactivate Proteinase K. Following centrifugation, the supernatant was used for analysis. PCRs were conducted with Phusion according to the manufacturer’s instructions (NEB).

To assess Arp53D RNA expression, whole flies (10 minimum) were ground in TRIzol (Invitrogen). Following centrifugation, the supernatant was chloroform extracted and the resulting soluble phase was isopropanol-extracted to precipitate RNA. RNA was then centrifuged, washed with 75% ethanol, dried and resuspended in RNAse-free water. Samples were treated with DNaseI (Zymo Research) or TURBO DNase (Thermo Fisher) according to the manufacturer’s instructions. DNase-treated samples were then further purified and concentrated using an RNA-cleanup kit (Zymo Research), and cDNA was obtained using SuperScript III first-strand synthesis (Invitrogen). All primers used are listed in Table S3.

### Immunoblot analysis

Approximately 30 testes from the transgenic line *w*^*-*^; *sfGFP-△N-term-Arp53D* and the line *w*^*-*^; *Arp53D KO; sfGFP-Arp53D* (full length) were dissected separately in PBS and centrifuged. After the supernatant was removed, the pellets of testes were flash frozen. Once thawed for immunoblot analysis, 20 µL of 4X NuPAGE LDS sample buffer (Thermo Fisher) was added to each pellet, which was resuspended and boiled for 5 min at 100°C. Protein samples were loaded on a mini-protean TGX stain-free protein gel (BioRad), run with Tris/Glycine/SDS buffer and transferred to a PVDF trans-blot turbo membrane (BioRad). After blocking with 5% milk in Tris-buffered saline (TBS) and 0.1% Tween-20 (TBST), the membrane was probed with anti-GFP and anti-tubulin in TBST for 1 hr at room temperature, followed by three 10 min washes with TBS. The membrane was then incubated for 45 min at room temperature with IR dye 680 anti-chicken (LI-COR) and/or IR dye 800 anti-rabbit 800 nm (LI-COR) in TBST (See Table S4 for dilutions). After 3 final washes with TBS, the membrane was scanned with 680 nm and 800 nm.

### Generation of the *Arp53D-KO* fly line

CRISPR/Cas9 was used to knockout *Arp53D* and replace it with *DsRed* to track the *Arp53D-KO* allele. Both guide RNAs were cloned into pCFD4 (Addgene 49411)^75^ and 1kb homology arms flanking *DsRed* were cloned into pHD-attP-DsRed (Addgene 51019)^76^. Guide RNAs were chosen based on optimal efficiency score and no predicted off-targets (http://www.flyrnai.org/crispr2/). The guide RNAs (TCCTGGAAACATGAGCAGCG and TTGGACGGGTGGTTCCGTCT) targeted internally to *Arp53D*, leading to an early stop-codon and removal of the actin fold domain. The CRISPR/Cas9 targets were chosen because they were least invasive to the nearby essential gene *SOD2* and predicted not to alter *SOD2’s* transcriptional regulatory elements. The two plasmids for CRISPR/Cas9 were midi-prepped (Takara Bio) and co-injected by BestGene, Inc in stock 55821 from the Bloomington *Drosophila* Stock Center (BDSC). BestGene, Inc. isolated transformants, crossed out the gene encoding for Cas9, and balanced the modified second chromosome with *CyO*. The *Arp53D-KO* fly line was backcrossed to the same Oregon-R fly line used in fertility assays for 8 generations, sequence verified and confirmed for lack of *Arp53D* expression and absence of *Wolbachia* (Figure S4C-G). The *Arp53D-KO* line was also separately backcrossed to the *w1118* fly line for 6 generations and sequence verified; this white-eyed line was subsequently used for cytological analyses and population cage experiments. For isogenization, females heterozygous for the *Arp53D-KO* allele were collected in each generation for a subsequent backcross since meiotic recombination only occurs in females, allowing for further mixing of genetic backgrounds. Heterozygous virgin flies were then crossed to obtain a homozygous *Arp53D-KO* fly strain, which was consistently maintained at room temperature and used for fertility assays.

### Fly culturing and generation of fly transgenics

All flies were cultured at 25°C on yeast-cornmeal-molasses-malt extract medium. *D. melanogaster Arp53D* was N-terminally tagged with *sfGFP* followed with a 6-aa intervening linker (GGSGGS). This transgene as well as all Arp53D variants *(△Nterm-Arp53D, Nterm Arp53D-Actin*, and *Nterm Arp53D-sfGFP*) were cloned into vectors encoding an attB site and *DsRed* under the control of an eye-specific promoter (3XP3). Constructs were midi-prepped (Takara Bio) and injected by BestGene, Inc. into BDSC 9744 (Table S5). To construct the *△N-term Arp53D* fly transgenic, 1-35 aa of Arp53D were removed. For the *Nterm Arp53D-Actin* fly line, 1-35 aa of Arp53D followed by the GGSGGS linker was added N-terminally to *D. melanogaster Act5C*. For *Nterm Arp53D-sfGFP*, Arp53D’s N-terminus (aa 1-35) followed by a 6-aa linker (GGSGGS) was added N-terminally to *sfGFP*, replacing Arp53D’s actin fold domain (aa 36-411); this construct did not have an N-terminal sfGFP tag. Transformants were selected, crossed to *w1118*, and were stably maintained as homozygous stocks. Modified sites were verified by PCR and subsequent Sanger sequencing.

### Immunofluorescence and live imaging

For live and fixed imaging, testes were dissected from 0-2-day old males in PBS using a dissecting scope. Live imaging was always conducted to confirm lack of fixation artifacts in immunofluorescence. For live imaging of individual cysts at all stages of spermatogenesis, dissected testes were transferred to a drop of PBS containing Hoechst 33342 (Invitrogen) and sir-actin (10 µM; Cytoskeleton, Inc.) on a slide and pulled apart, evenly distributing visibly elongated cysts. Cysts were stained for 5 min at room temperature, and then a coverslip was placed on top for imaging.

To image whole fixed testes, dissected testes were immediately fixed with 2% PFA in periodate-lysine-paraformaldehyde (PLP) buffer for 1 hr at room temperature, and then permeabilized with PBS with 0.5% Triton X-100 for 30 min. Testes were blocked for 30 min with 3% BSA in PBS plus 0.1% Triton X-100 (PBST). Incubation with primary antibodies took place overnight at 4°C. Testes were then washed several times, followed by secondary antibody incubation for 2 hr at room temperature. After washing 3 times with PBST, testes were mounted onto slides with VECTASHIELD antifade mounting media with DAPI (Thermo Fisher).

For fixation of individual cysts, cysts were separated (as done for live imaging) in PBS without Hoechst. After a coverslip was placed on the slide, it was submerged in liquid nitrogen. Tissue was fixed with either paraformaldehyde (PFA) or methanol. For PFA fixation, the coverslip was removed from flash-frozen slides and the slides were placed in 100% ethanol for 10 min. Then fixation with 4% PFA in PBS took place for 7 min at room temperature. Tissue was then permeabilized twice for 15 min each with PBS and 0.3% Triton X-100 and 0.3% sodium deoxycholate. Alternative fixation with methanol took place at -20°C for 5 min, followed by incubation in acetone -20°C for 5 min. After both fixation protocols, slides were washed once with PBST for 10 min and then blocked with 3% BSA in PBST for 30 min. Primary antibody incubations took place overnight at 4°C, followed by three 15 min washes in PBS at room temperature. Slides were incubated with secondary antibody for 1 hr, followed by four 15-min washes with PBS. Slides were then either washed once with Hoechst 33342 (Invitrogen) or DNA-stained with mounting media containing DAPI (Thermo Fisher). Following the addition of mounting media, a coverslip was placed and sealed with nail polish. See Table S4 for antibody dilutions. All live and fixed samples were imaged using a confocal microscope (Leica TCS SP5 II) and LASAF software (Leica).

### Fertility assays

Oregon-R flies were used as wild type flies, and female and male virgins were collected for all assays. Females were 1-5 days old, and males were 1-2 days old. Crosses were setup with females 4-fold in excess, and matings took place over 9 days at 25°C or 29°C with vials flipped every 2-3 days. Light/dark cycles were maintained consistently. For heat-stressed experiments, virgins were maintained at 25°C until crosses were setup and then transferred to 29°C. All adult progeny were quantified on the last possible day before the second-generation progeny emerged. With day 1 being the time at which crosses were set up, day 15 or 16 was the last day the first generation could be counted at 25°C, and for 29°C experiments, day 12 or 13 was the last day. For quantification of fluorescence (the *Arp53D-KO* allele) in the Oregon-R background, DsRed fluorescence was visualized in the ocelli because pigmentation obscured fluorescence in the eye. Heterozygous flies were generated by crossing KO females to Oregon-R flies at room temperature. Homozygous *DsRed* flies were denoted by strikingly fluorescent ocelli and dim fluorescence of the body, whereas the ocelli of heterozygous flies were dim and required close observation to differentiate from wild-type flies. All fertility assays were conducted at a minimum of n=2, and parents that died in all crosses were tallied with each passage and did not differ significantly among the different genotypes.

To compare development of embryos, female and male virgins were collected as done for the previously described fertility assays. Fly crosses were set up with females in 4-fold excess and allowed to lay at 29°C for 2 hr. The number of eggs laid were quantified and then returned to 29°C. After 24 hr, unhatched embryos were quantified.

For knockdown of *Arp53D*, RNAi line 108369 (VDRC) was used and sequence verified (as done in Green et al., 2014) for integration at the chromosomal 30B site and not the 40D site, which has a non-specific phenotype^77^. The line was crossed to *topi-Gal4* flies (generously given by the labs of Lynn Cooley and Christian Lehner) for knockdown in late spermatogenesis.

### Population cage experiment

The isogenized *Arp53D-KO* line in the *w1118* background was used due to ease of DsRed detection in the eye (as opposed to the ocelli in the Oregon-R *Arp53D-KO* background). Virgin females and males were collected from the *w1118* fly line and the *Arp53D-KO* fly line isogenized in the *w1118* background. Crosses with 50 *Arp53D-KO* females, 25 *Arp53D-KO* males and 25 *w1118* males were set up in bottles with 3 replicates. Crosses were passaged every 2 weeks at room temperature. At each passage, 50 females and 50 males were randomly collected without fluorescence detection and without selection based on virgin status. The 100 progeny were placed in a fresh bottle and remaining progeny were frozen for subsequent detection of DsRed fluorescence. After one week, parents of the next generation were removed and frozen to include in the previous generation’s quantification.

## Acknowledgements

We thank Janet Young and Grace Yuh Chwen for help with the McDonald-Kreitman analyses and Ching-Ho Chang for help with RNA-seq analysis. We thank Susan Parkhurst and Barbara Wakimoto for providing fly lines and technical advice and Akhila Rajan for lending the critical 29°C room used for fertility experiments. We also thank Mollie Manier for offering dissection protocols and helpful discussions about *Arp53D-KO* phenotypes, Leslie Vosshall for providing anti-Androcam antibodies, and Kathleen Beckingham for discussing potential co-localization of Androcam and Arp53D. Lastly, we thank Ching-Ho Chang, Lisa Kursel, Antoine Molaro, and Pravrutha Raman for providing comments on the manuscript as well as the rest of the Malik lab for useful discussions and *Drosophila* training. This work was funded by the Jane Coffin Childs Memorial Fund (CMS), an NIGMS K99 Pathway to Independence Award (CMS), NIH grant R01GM074108 (HSM) and the Howard Hughes Medical Institute (HSM).

## References

1. Dominguez, R. & Holmes, K.C. Actin structure and function. Annu Rev Biophys 40, 169–86 (2011).

2. Schrank, B.R. et al. Nuclear ARP2/3 drives DNA break clustering for homology-directed repair. Nature 559, 61–66 (2018).

3. Wei, M. et al. Nuclear actin regulates inducible transcription by enhancing RNA polymerase II clustering. Science 6(2020).

4. Goodson, H.V. & Hawse, W.F. Molecular evolution of the actin family. J Cell Sci 115, 2619–22 (2002).

5. Muller, J. et al. Sequence and comparative genomic analysis of actin-related proteins. Mol Biol Cell 16, 5736–48 (2005).

6. van den Ent, F., Amos, L.A. & Lowe, J. Prokaryotic origin of the actin cytoskeleton. Nature 413, 39–44 (2001).

7. Izore, T., Kureisaite-Ciziene, D., McLaughlin, S.H. & Lowe, J. Crenactin forms actin-like double helical filaments regulated by arcadin-2. Elife 5(2016).

8. Mullins, R.D., Heuser, J.A. & Pollard, T.D. The interaction of Arp2/3 complex with actin: nucleation, high affinity pointed end capping, and formation of branching networks of filaments. Proc Natl Acad Sci U S A 95, 6181–6 (1998).

9. Harata, M. et al. Multiple actin-related proteins of Saccharomyces cerevisiae are present in the nucleus. J Biochem 128, 665–71 (2000).

10. Blessing, C.A., Ugrinova, G.T. & Goodson, H.V. Actin and ARPs: action in the nucleus. Trends Cell Biol 14, 435–42 (2004).

11. Klages-Mundt, N.L., Kumar, A., Zhang, Y., Kapoor, P. & Shen, X. The Nature of Actin-Family Proteins in Chromatin-Modifying Complexes. Front Genet 9, 398 (2018).

12. Muhua, L., Karpova, T.S. & Cooper, J.A. A yeast actin-related protein homologous to that in vertebrate dynactin complex is important for spindle orientation and nuclear migration. Cell 78, 669–79 (1994).

13. Lee, I.H., Kumar, S. & Plamann, M. Null mutants of the neurospora actin-related protein 1 pointed-end complex show distinct phenotypes. Mol Biol Cell 12, 2195–206 (2001).

14. Liu, S.L., May, J.R., Helgeson, L.A. & Nolen, B.J. Insertions within the actin core of actin-related protein 3 (Arp3) modulate branching nucleation by Arp2/3 complex. J Biol Chem 288, 487–97 (2013).

15. Chen, M. & Shen, X. Nuclear actin and actin-related proteins in chromatin dynamics. Curr Opin Cell Biol 19, 326–30 (2007).

16. Machesky, L.M. & May, R.C. Arps: actin-related proteins. Results Probl Cell Differ 32, 213–29 (2001).

17. Fyrberg, C., Ryan, L., Kenton, M. & Fyrberg, E. Genes encoding actin-related proteins of Drosophila melanogaster. J Mol Biol 241, 498–503 (1994).

18. Schroeder, C.M., Valenzuela, J.R., Natividad, I.M., Hocky, G.M. & Malik, H.S. A burst of genetic innovation in Drosophila actin-related proteins for testis-specific function. Mol Biol Evol (2019).

19. Heid, H. et al. Novel actin-related proteins Arp-T1 and Arp-T2 as components of the cytoskeletal calyx of the mammalian sperm head. Exp Cell Res 279, 177–87 (2002).

20. Tanaka, H. et al. Novel actin-like proteins T-ACTIN 1 and T-ACTIN 2 are differentially expressed in the cytoplasm and nucleus of mouse haploid germ cells. Biol Reprod 69, 475–82 (2003).

21. Hara, Y., Yamagata, K., Oguchi, K. & Baba, T. Nuclear localization of profilin III-ArpM1 complex in mouse spermiogenesis. FEBS Lett 582, 2998–3004 (2008).

22. Boeda, B. et al. Molecular recognition of the Tes LIM2-3 domains by the actin-related protein Arp7A. J Biol Chem 286, 11543–54 (2011).

23. Fu, J. et al. Anti-ACTL7a antibodies: a cause of infertility. Fertil Steril 97, 1226-33 e1-8 (2012).

24. Drosophila 12 Genomes, C. Evolution of genes and genomes on the Drosophila phylogeny. Nature 450, 203–18 (2007).

25. Chen, Z.X. et al. Comparative validation of the D. melanogaster modENCODE transcriptome annotation. Genome Res 24, 1209–23 (2014).

26. Zhou, Q. & Bachtrog, D. Sex-specific adaptation drives early sex chromosome evolution in Drosophila. Science 337, 341–5 (2012).

27. Renschler, G. et al. Hi-C guided assemblies reveal conserved regulatory topologies on X and autosomes despite extensive genome shuffling. Genes Dev 33, 1591–1612 (2019).

28. Kurek, R., Reugels, A.M., Glatzer, K.H. & Bunemann, H. The Y chromosomal fertility factor Threads in Drosophila hydei harbors a functional gene encoding an axonemal dynein beta heavy chain protein. Genetics 149, 1363–76 (1998).

29. Lynch, M. & Conery, J.S. The evolutionary fate and consequences of duplicate genes. Science 290, 1151–5 (2000).

30. Lack, J.B. et al. The Drosophila genome nexus: a population genomic resource of 623 Drosophila melanogaster genomes, including 197 from a single ancestral range population. Genetics 199, 1229–41 (2015).

31. Lack, J.B., Lange, J.D., Tang, A.D., Corbett-Detig, R.B. & Pool, J.E. A Thousand Fly Genomes: An Expanded Drosophila Genome Nexus. Mol Biol Evol 33, 3308–3313 (2016).

32. Hervas, S., Sanz, E., Casillas, S., Pool, J.E. & Barbadilla, A. PopFly: the Drosophila population genomics browser. Bioinformatics 33, 2779–2780 (2017).

33. McDonald, J.H. & Kreitman, M. Adaptive protein evolution at the Adh locus in Drosophila. Nature 351, 652–4 (1991).

34. Fay, J.C., Wyckoff, G.J. & Wu, C.I. Positive and negative selection on the human genome. Genetics 158, 1227–34 (2001).

35. Bierne, N. & Eyre-Walker, A. The genomic rate of adaptive amino acid substitution in Drosophila. Mol Biol Evol 21, 1350–60 (2004).

36. Celniker, S.E. et al. Unlocking the secrets of the genome. Nature 459, 927–30 (2009).

37. Fabian, L. & Brill, J.A. Drosophila spermiogenesis: Big things come from little packages. Spermatogenesis 2, 197–212 (2012).

38. de Cuevas, M. & Spradling, A.C. Morphogenesis of the Drosophila fusome and its implications for oocyte specification. Development 125, 2781–9 (1998).

39. Lin, H., Yue, L. & Spradling, A.C. The Drosophila fusome, a germline-specific organelle, contains membrane skeletal proteins and functions in cyst formation. Development 120, 947–56 (1994).

40. de Cuevas, M., Lee, J.K. & Spradling, A.C. alpha-spectrin is required for germline cell division and differentiation in the Drosophila ovary. Development 122, 3959–68 (1996).

41. Noguchi, T. & Miller, K.G. A role for actin dynamics in individualization during spermatogenesis in Drosophila melanogaster. Development 130, 1805–16 (2003).

42. Fabrizio, J.J., Hime, G., Lemmon, S.K. & Bazinet, C. Genetic dissection of sperm individualization in Drosophila melanogaster. Development 125, 1833–43 (1998).

43. Noguchi, T., Lenartowska, M., Rogat, A.D., Frank, D.J. & Miller, K.G. Proper cellular reorganization during Drosophila spermatid individualization depends on actin structures composed of two domains, bundles and meshwork, that are differentially regulated and have different functions. Mol Biol Cell 19, 2363–72 (2008).

44. Rogat, A.D. & Miller, K.G. A role for myosin VI in actin dynamics at sites of membrane remodeling during Drosophila spermatogenesis. J Cell Sci 115, 4855–65 (2002).

45. Frank, D.J. et al. Androcam is a tissue-specific light chain for myosin VI in the Drosophila testis. J Biol Chem 281, 24728–36 (2006).

46. Wasbrough, E.R. et al. The Drosophila melanogaster sperm proteome-II (DmSP-II). J Proteomics 73, 2171–85 (2010).

47. Serbus, L.R., Casper-Lindley, C., Landmann, F. & Sullivan, W. The genetics and cell biology of Wolbachia-host interactions. Annu Rev Genet 42, 683–707 (2008).

48. Raychaudhuri, N. et al. Transgenerational propagation and quantitative maintenance of paternal centromeres depends on Cid/Cenp-A presence in Drosophila sperm. PLoS Biol 10, e1001434 (2012).

49. Jevitt, A. et al. A single-cell atlas of adult Drosophila ovary identifies transcriptional programs and somatic cell lineage regulating oogenesis. PLoS Biol 18, e3000538 (2020).

50. Slaidina, M., Banisch, T.U., Gupta, S. & Lehmann, R. A single-cell atlas of the developing Drosophila ovary identifies follicle stem cell progenitors. Genes Dev 34, 239–249 (2020).

51. Jambor, H. et al. Systematic imaging reveals features and changing localization of mRNAs in Drosophila development. Elife 4(2015).

52. Karaiskos, N. et al. The Drosophila embryo at single-cell transcriptome resolution. Science 358, 194–199 (2017).

53. Lott, S.E. et al. Noncanonical compensation of zygotic X transcription in early Drosophila melanogaster development revealed through single-embryo RNA-seq. PLoS Biol 9, e1000590 (2011).

54. Sutoh, K., Ando, M., Sutoh, K. & Toyoshima, Y.Y. Site-directed mutations of Dictyostelium actin: disruption of a negative charge cluster at the N terminus. Proc Natl Acad Sci U S A 88, 7711–4 (1991).

55. Hansen, J.E., Marner, J., Pavlov, D., Rubenstein, P.A. & Reisler, E. Structural transition at actin’s N-terminus in the actomyosin cross-bridge cycle. Biochemistry 39, 1792–9 (2000).

56. Varland, S., Vandekerckhove, J. & Drazic, A. Actin Post-translational Modifications: The Cinderella of Cytoskeletal Control. Trends Biochem Sci 44, 502–516 (2019).

57. de Cuevas, M., Lilly, M.A. & Spradling, A.C. Germline cyst formation in Drosophila. Annu Rev Genet 31, 405–28 (1997).

58. Kloc, M., Bilinski, S., Dougherty, M.T., Brey, E.M. & Etkin, L.D. Formation, architecture and polarity of female germline cyst in Xenopus. Dev Biol 266, 43–61 (2004).

59. Geyer, C.B. et al. A missense mutation in the Capza3 gene and disruption of F-actin organization in spermatids of repro32 infertile male mice. Dev Biol 330, 142–52 (2009).

60. Kleene, K.C. Sexual selection, genetic conflict, selfish genes, and the atypical patterns of gene expression in spermatogenic cells. Dev Biol 277, 16–26 (2005).

61. Swanson, W.J. & Vacquier, V.D. The rapid evolution of reproductive proteins. Nat Rev Genet 3, 137–44 (2002).

62. Panhuis, T.M., Clark, N.L. & Swanson, W.J. Rapid evolution of reproductive proteins in abalone and Drosophila. Philos Trans R Soc Lond B Biol Sci 361, 261–8 (2006).

63. Figard, L. et al. Cofilin-Mediated Actin Stress Response Is Maladaptive in Heat-Stressed Embryos. Cell Rep 26, 3493–3501 e4 (2019).

64. Vinckenbosch, N., Dupanloup, I. & Kaessmann, H. Evolutionary fate of retroposed gene copies in the human genome. Proc Natl Acad Sci U S A 103, 3220–5 (2006).

65. Nyberg, K.G. & Carthew, R.W. Out of the testis: biological impacts of new genes. Genes Dev 31, 1825–1826 (2017).

66. Kursel, L.E. & Malik, H.S. Gametic specialization of centromeric histone paralogs in Drosophila virilis. bioRxiv (2019).

67. Ross, B.D. et al. Stepwise evolution of essential centromere function in a Drosophila neogene. Science 340, 1211–4 (2013).

68. Chebbo, S., Josway, S., Belote, J.M. & Manier, M.K. A putative novel role for Eip74EF in male reproduction in promoting sperm elongation at the cost of male fecundity. J Exp Zool B Mol Dev Evol (2020).

69. VanKuren, N.W. & Long, M. Gene duplicates resolving sexual conflict rapidly evolved essential gametogenesis functions. Nat Ecol Evol 2, 705–712 (2018).

70. Thurmond, J. et al. FlyBase 2.0: the next generation. Nucleic Acids Res 47, D759–D765 (2019).

71. Katoh, K. & Standley, D.M. MAFFT multiple sequence alignment software version 7: improvements in performance and usability. Mol Biol Evol 30, 772–80 (2013).

72. Kearse, M. et al. Geneious Basic: an integrated and extendable desktop software platform for the organization and analysis of sequence data. Bioinformatics 28, 1647–9 (2012).

73. Egea, R., Casillas, S. & Barbadilla, A. Standard and generalized McDonald-Kreitman test: a website to detect selection by comparing different classes of DNA sites. Nucleic Acids Res 36, W157–62 (2008).

74. Hu, T.T., Eisen, M.B., Thornton, K.R. & Andolfatto, P. A second-generation assembly of the Drosophila simulans genome provides new insights into patterns of lineage-specific divergence. Genome Res 23, 89–98 (2013).

75. Port, F., Chen, H.M., Lee, T. & Bullock, S.L. Optimized CRISPR/Cas tools for efficient germline and somatic genome engineering in Drosophila. Proc Natl Acad Sci U S A 111, E2967–76 (2014).

76. Gratz, S.J. et al. Highly specific and efficient CRISPR/Cas9-catalyzed homology-directed repair in Drosophila. Genetics 196, 961–71 (2014).

77. Green, E.W., Fedele, G., Giorgini, F. & Kyriacou, C.P. A Drosophila RNAi collection is subject to dominant phenotypic effects. Nat Methods 11, 222–3 (2014).

78. Ishida, T. & Kinoshita, K. PrDOS: prediction of disordered protein regions from amino acid sequence. Nucleic Acids Res 35, W460–4 (2007).

79. Schneider, D.I., Klasson, L., Lind, A.E. & Miller, W.J. More than fishing in the dark: PCR of a dispersed sequence produces simple but ultrasensitive Wolbachia detection. BMC Microbiol 14, 121 (2014).

80. Kent, W.J. BLAT--the BLAST-like alignment tool. Genome Res 12, 656–64 (2002).

81. Venken, K.J., He, Y., Hoskins, R.A. & Bellen, H.J. P[acman]: a BAC transgenic platform for targeted insertion of large DNA fragments in D. melanogaster. Science 314, 1747–51 (2006).

