## Supplementary Figures and Tables for "A rapidly evolving actin mediates fertility and developmental tradeoffs in *Drosophila*"

**A**

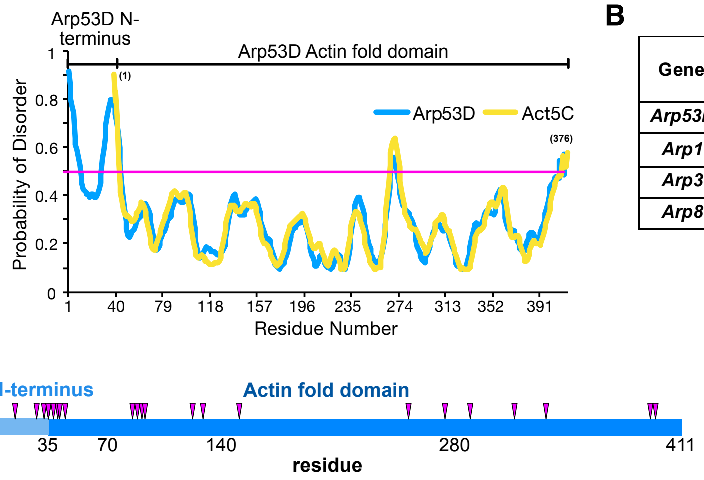


**B**

| Gene | Polymorphisms | | Divergences | | Unpolarized | |
|---|---|---|---|---|---|---|
| | $P_S$ | $P_N$ | $D_S$ | $D_N$ | p-value | α |
| ***Arp53D*** | 19 | 1 | 24 | 21 | **0.001** | **0.939** |
| ***Arp1*** | 48 | 2 | 19 | 1 | 1 | 0.21 |
| ***Arp3*** | 34 | 1 | 16 | 0 | 1 | NA |
| ***Arp8*** | 96 | 30 | 27 | 7 | 0.82 | -0.21 |

**D**

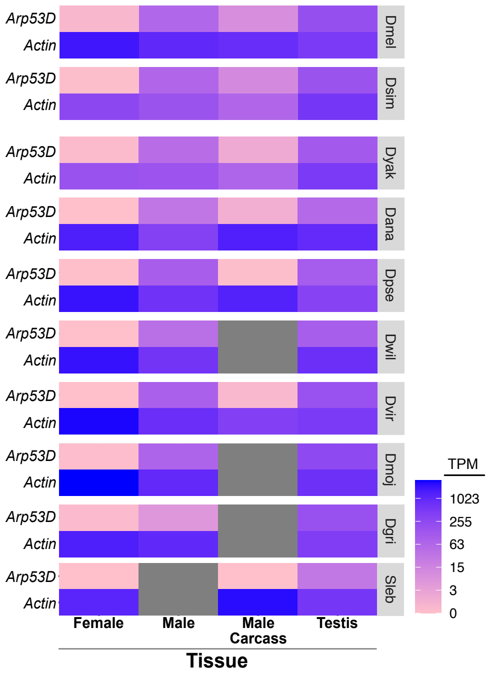


**Supplementary Figure S1. Arp53D diverged in sequence and expression from actin. A)** The probability of disorder of the protein sequence of *D. melanogaster* Arp53D (blue) and Act5C (yellow) are graphed per amino acid. Probability was determined using PrDOS[78]. **B)** *Arp53D* and canonical Arps (Arp1, 3 and Arp8) from 197 *D. melanogaster* alleles (DPGP3)[30] were aligned to their orthologous sequences in *D. simulans* [74] (outgroup) in an unpolarized MKT test, which compares the ratio of nonsynonymous to synonymous polymorphisms ($P_N$ and $P_S$) among the *D. melanogaster* strains and the ratio of nonsynonymous to synonymous fixed substitutions ($D_N$ and $D_S$) between the *D. melanogaster* and *D. simulans* orthologs. The alignment was manually checked with Clustal Omega and rare variants in Arp53D (<5% of sequences) were removed. Positive selection (blue, bold text) is indicated with an α-value approaching 1 and a statistically significant p-value. Canonical Arps 1, 3 and 8 showed no positive selection with and without rare variants removed. **C)** Non-synonymous fixed changes ($D_N$) identified by the unpolarized McDonald-Kreitman test are located in the schematic representing Arp53D's protein sequence (411 aa). **D)** RNA-seq data (levels in TPM) from females, males, male carcass, and the testis are displayed for 10 *Drosophila* species with lowest values shown in pink and highest values in blue. See Table S2 for accession numbers of the datasets used.

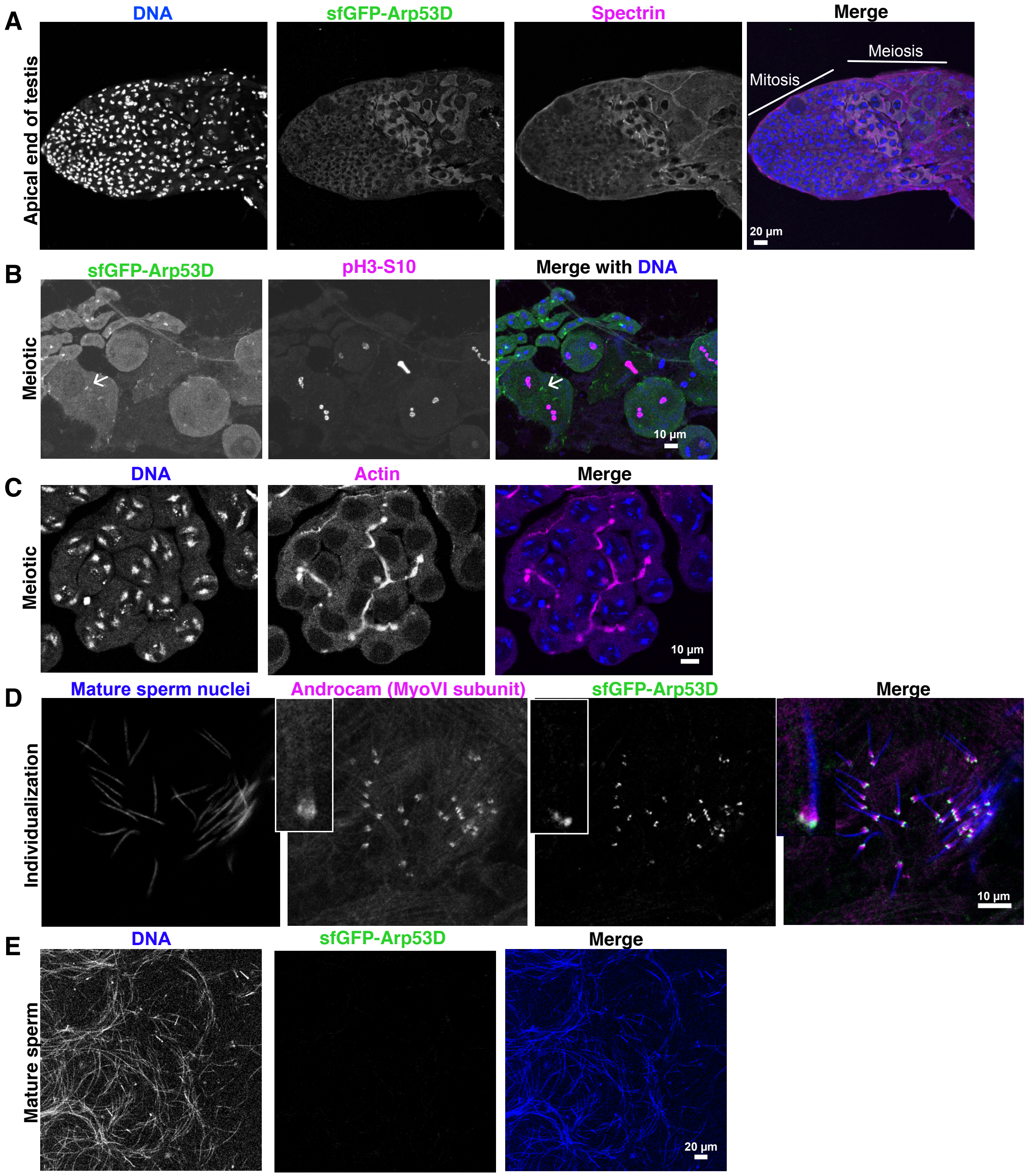


**Supplementary Figure S2. Arp53D is expressed in meiosis and co-localizes with a testis-specific myosin VI subunit. A)** Testes expressing sfGFP-Arp53D were fixed and probed with anti-GFP and anti-α-spectrin.GFP signal is detected in meiotic cells and not mitotic cells at the apical end of the testis. **B)** Testes expressing sfGFP-Arp53D were fixed and probed with anti-GFP and anti-phosphorylated (S10) histone H3 (pH3-S10), a marker for meiosis. sfGFP-Arp53D (indicated by arrow) is detected in cells that are meiotic (positive for pH3-S10). **C)** Testis meiotic cells were probed for DNA and actin (phalloidin probe) to show actin localizes to the male fusome. **D)** Testes expressing sfGFP-Arp53D were fixed and probed with anti-GFP and anti-Androcam, the testis-specific light chain of myosin VI[45]. Arp53D and Androcam localize at the leading edge of actin cones at mature sperm nuclei, shown in blue (DAPI). **E)** No GFP signal was detected in mature sperm (denoted by condensed DNA, blue) dissected from the seminal vesicle of sfGFP-Arp53D-expressing flies.

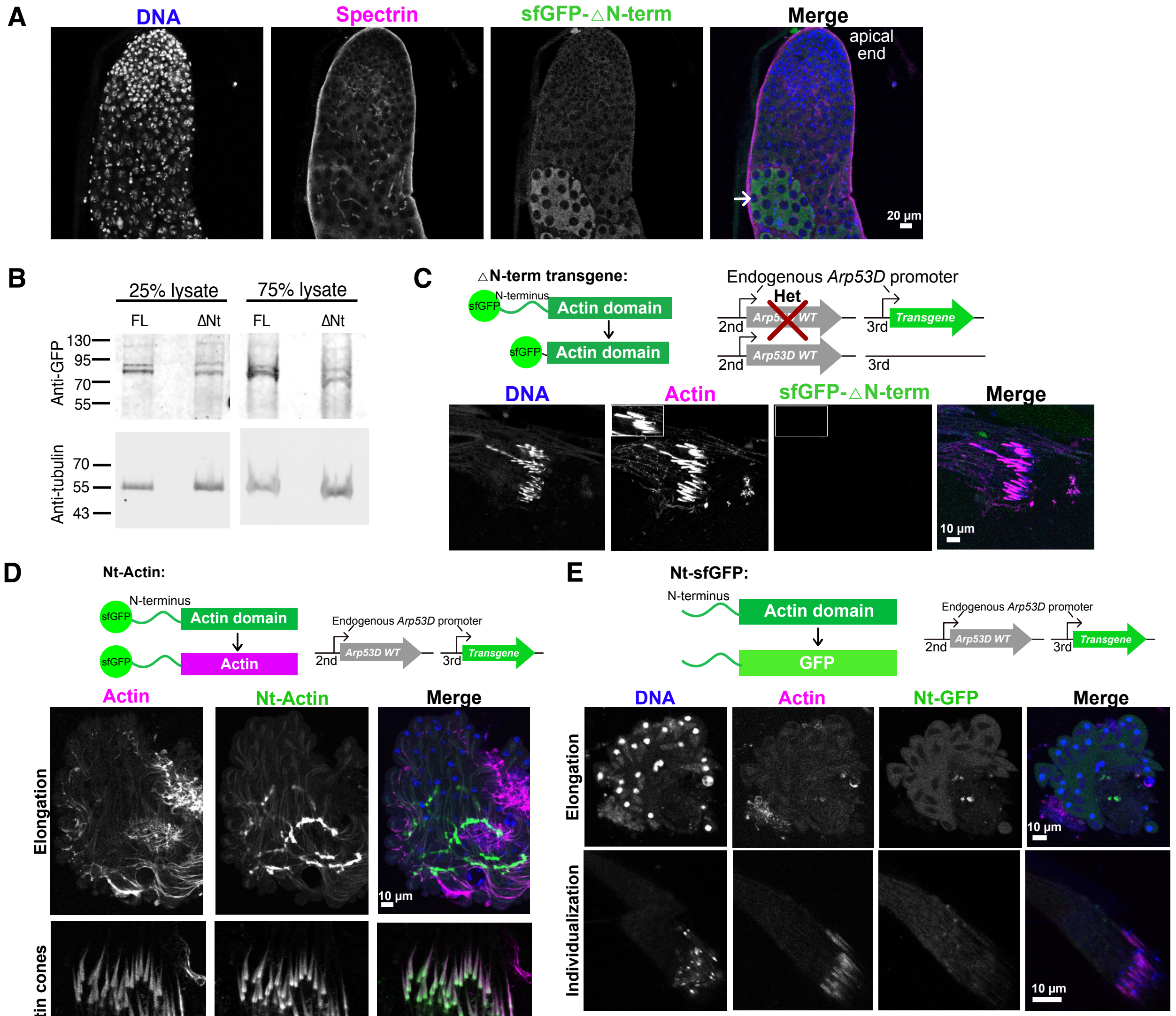

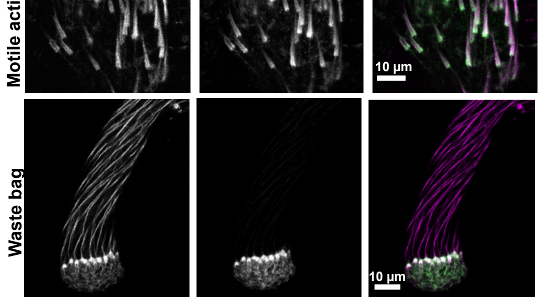


**Supplementary Figure S3. Chimeric proteins reveal Arp53D's unique N-terminus is required for localization to germline actin structures. A)** Whole testes expressing sfGFP-△N-term were dissected, fixed and probed with anti-spectrin (magenta), anti-GFP (green) and DAPI (blue). GFP is detected in meiosis (indicated by arrow). **B)** The expression of sfGFP-△N-term Arp53D was compared to that of sfGFP-tagged full-length Arp53D by denaturing 15 pairs of dissected testes from each fly line and conducting SDS-PAGE analysis. Immunoblots were probed with anti-GFP and anti-tubulin (loading control). Amounts loaded are displayed as a percentage of total lysate. **C)** A transgene on the third chromosome encoded "△N-term", sfGFP-Arp53D with 35 aa of the N-terminus removed. Homozygotes of the transgenic line were crossed to Arp53D-KOs, creating flies heterozygous for *Arp53D-KO* and *sfGFP-△N-term*. Testes were dissected and imaged live. No GFP signal co-localized with actin cones (sir-actin probe). **D)** Testes from the transgenic fly line expressing a chimeric form of Arp53D (actin with Arp53D's N-terminus, "Nt-Actin") were dissected and imaged live. GFP localizes to the fusome of elongating spermatids, motile actin cones (no longer co-localizing with mature sperm nuclei) and the waste bag. **E)** The actin domain of Arp53D was replaced with sfGFP, and the transgene ("Nt-sfGFP") was targeted to the third chromosome. Testes were dissected and imaged live with Hoechst (DNA, blue) and sir-actin (actin, magenta). No GFP signal co-localized with the fusome in elongating cells or to actin cones during individualization.

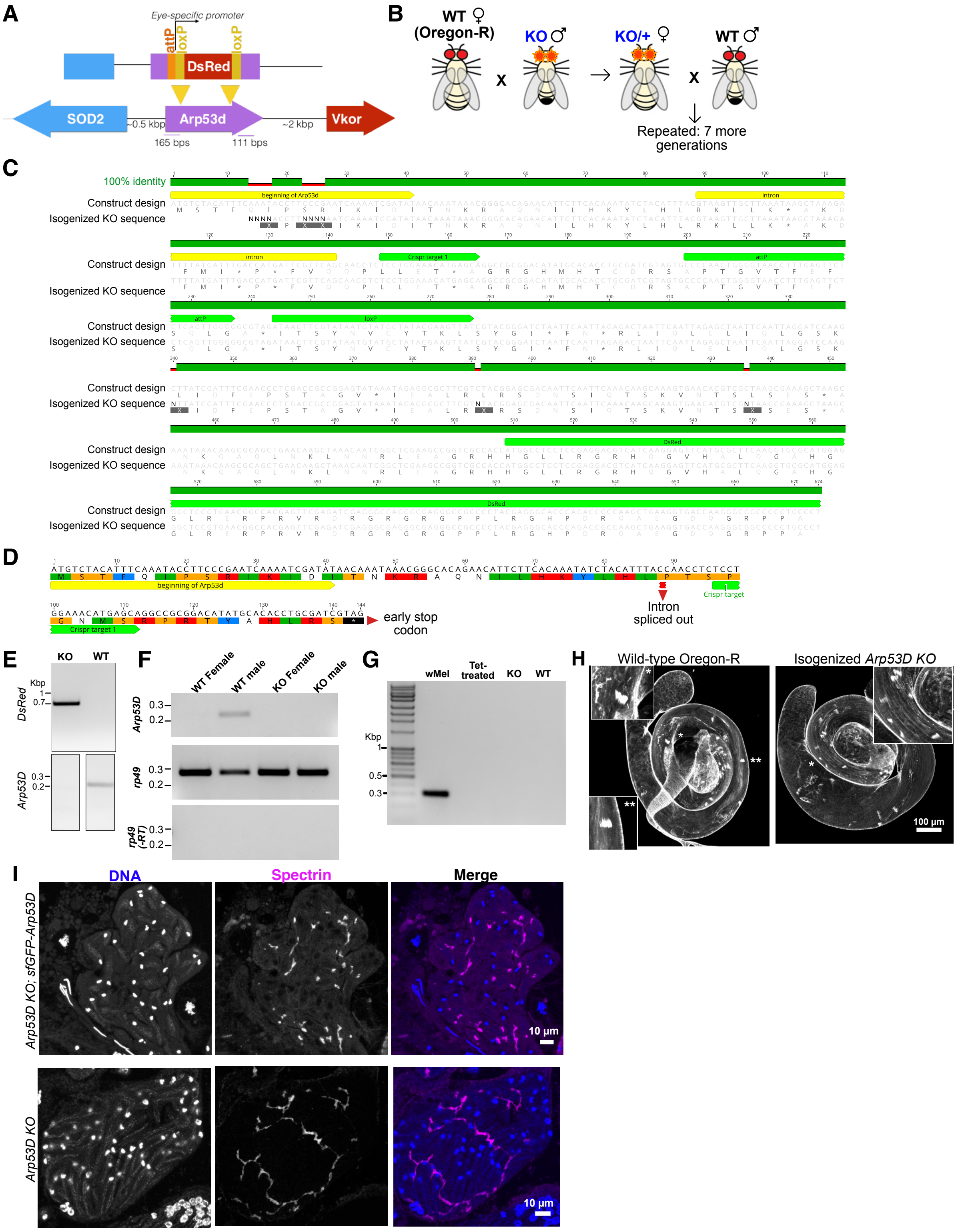


**Supplementary Figure S4. Characterization of isogenized *Arp53D-KO* flies verifies CRISPR-Cas9 deletion. A)** The Arp53D gene was targeted using CRISPR/Cas9 with PAM sites internal to Arp53D. A gene encoding for the fluorescent protein DsRed was knocked-into the locus, creating an early stop codon. DsRed is under the control of an eye-specific promoter (3xP3) and is flanked by an attP site and loxP sites. **B)** The Arp53D knockout (KO) was backcrossed to the wildtype Oregon-R *D. melanogaster strain*. Male KOs were crossed to

**Supplementary Figure S4, continued**

female Oregon-R, and female progeny heterozygous for the KO allele were collected and crossed to Oregon-R males. Heterozygous females were selected because unlike males, meiotic recombination takes place and facilitates isogenization. Backcrosses were repeated for a total of 8 generations. **C)** The *Arp53D-KO* locus (PCR product in panel E) was sequenced and aligned with the designed sequence. Annotations indicate one CRISPR target site and the attP, loxP and DsRed sites in the knock-in construct. **D)** The CRISPR/Cas9 cut sites yielded an early stop codon with only 37 aa of Arp53D remaining. **E)** To sequence-verify the *Arp53D-KO* genomic locus, a PCR was conducted with genomic DNA and primers upstream from *Arp53D* and within the gene encoding *DsRed*, yielding a ~700 bp product. The WT line used indicated no presence of *DsRed* in the *Arp53D* locus ("DsRed," top row). A PCR with primers targeting the actin fold domain of *Arp53D* showed a band in the WT line and not the KO as expected ("Arp53D," bottom row). **F)** RNA was extracted from Oregon-R and isogenized *Arp53D-KO* females and males. RNA was reverse transcribed and RT-PCRs (25 cycles) were done to assess expression of *Arp53D*. *Rp49* was used to compare amounts between samples, and template without reverse transcriptase was used to verify absence of genomic DNA. **G)** The Oregon-R and isogenized *Arp53D-KO* lines were tested for presence of *Wolbachia* with primers targeting the ARM locus[79]. The KO, WT and tetracycline-treated Bloomington *Drosophila* Stock Center (BDSC) line #145 ("Tet-treated") yielded no PCR product unlike *Wolbachia*-infected BDSC #145 ("wMel"), which yielded the expected 300-bp amplicon. **H)** Oregon-R and isogenized *Arp53D-KO* whole testes were fixed and actin was labeled with sir-actin. Insets show enlarged actin cones and asterisks indicate their locations in the testis. **I)** Elongating cysts from *Arp53D-KO* and *Arp53D-KO; sfGFP-Arp53D* were dissected and stained with DAPI (DNA) and anti-spectrin (fusome).

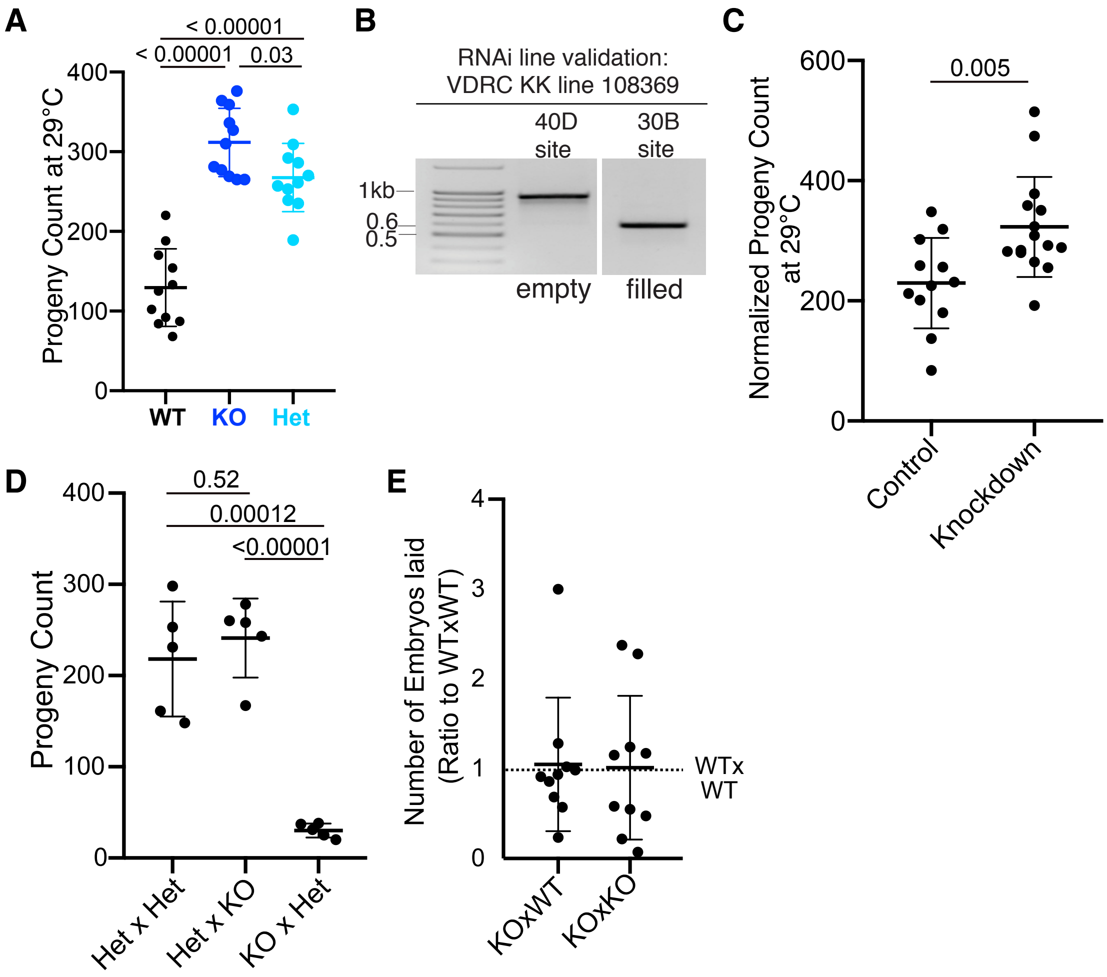


**Supplementary Figure S5. Male and female fertility are impacted by lack of Arp53D. A)** A fertility assay was conducted at 29°C with WT females crossed to WT, KO or heterozygous ("Het") *Arp53D* males. Adult progeny were counted. **B)** Targeted sequencing of two possible integration sites in the VDRC RNAi KK line 108369 was done and indicated integration of the RNAi hairpin at chromosomal site 30B and not 40D. The 30B site is preferable, as the 40D site gives non-specific phenotypes[77]. **C)** A fertility assay at 29°C was conducted with *Arp53D*-knockdown males. The male knockdown flies encoded topi-Gal4, which is expressed in late spermatogenesis, and an RNAi hairpin targeting *Arp53D* (VDRC KK line 108369 in B). The control male encoded only the RNAi hairpin. Mating and embryo laying took place over 9 days, and adult progeny were counted and normalized by accounting for number of dead parents: total progeny * total parents/(total parents - dead parents). **D)** Het and KO females and males (female x male) were crossed at 29°C for a week and total adult progeny were counted. All p-values were determined with a one-way ANOVA test. **E)** WTxWT, KOxWT and KOxKO were allowed to lay for the same amount of time at 29°C, and the number of embryos laid were quantified immediately following mating. Data was normalized to WTxWT and is shown as a ratio, with the ratio of 1 for WTxWT indicated by a dotted line.

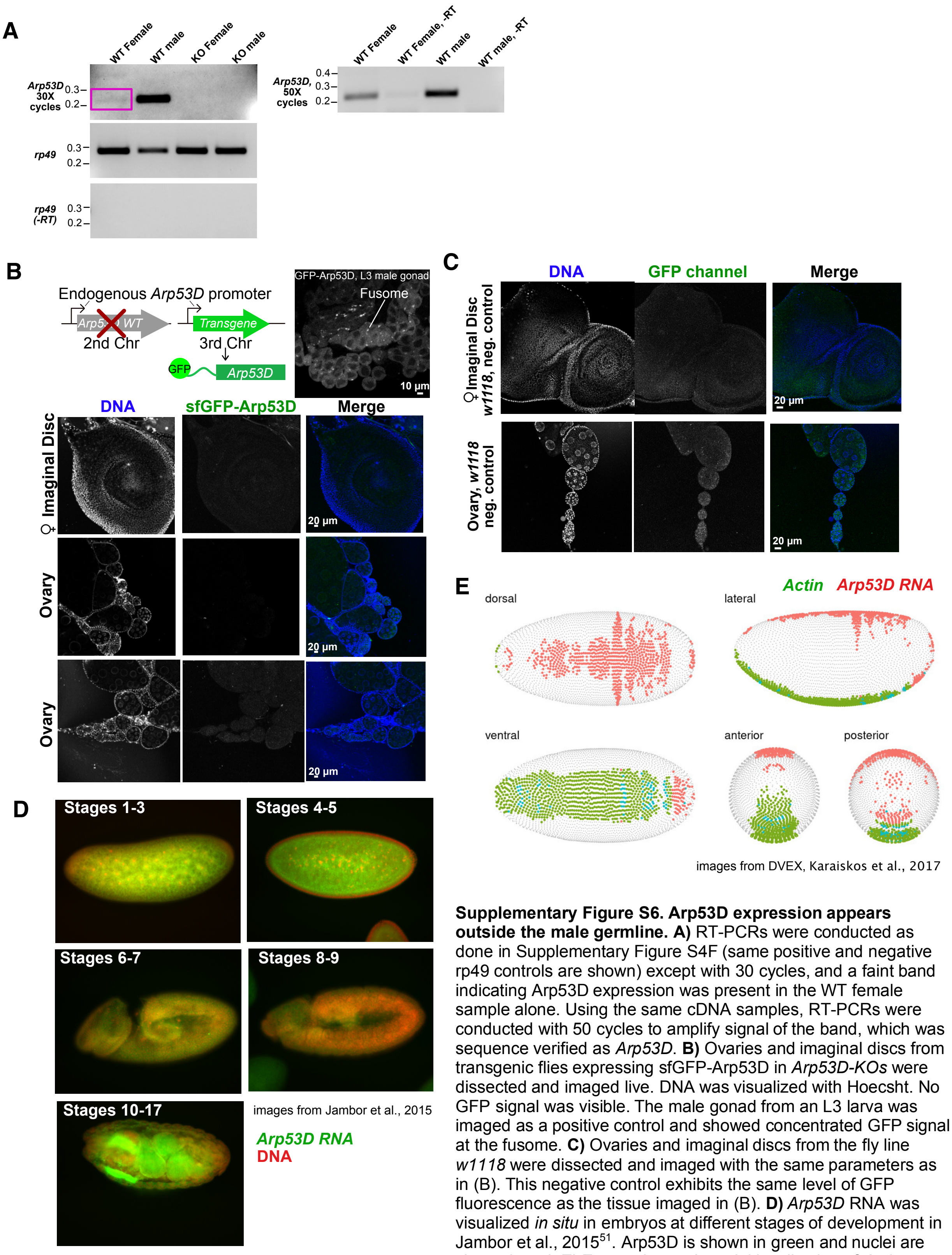


**Supplementary Figure S6. Arp53D expression appears outside the male germline. A)** RT-PCRs were conducted as done in Supplementary Figure S4F (same positive and negative rp49 controls are shown) except with 30 cycles, and a faint band indicating Arp53D expression was present in the WT female sample alone. Using the same cDNA samples, RT-PCRs were conducted with 50 cycles to amplify signal of the band, which was sequence verified as *Arp53D*. **B)** Ovaries and imaginal discs from transgenic flies expressing sfGFP-Arp53D in *Arp53D-KOs* were dissected and imaged live. DNA was visualized with Hoecsht. No GFP signal was visible. The male gonad from an L3 larva was imaged as a positive control and showed concentrated GFP signal at the fusome. **C)** Ovaries and imaginal discs from the fly line *w1118* were dissected and imaged with the same parameters as in (B). This negative control exhibits the same level of GFP fluorescence as the tissue imaged in (B). **D)** *Arp53D* RNA was visualized *in situ* in embryos at different stages of development in Jambor et al., 2015[51]. Arp53D is shown in green and nuclei are shown in red. **E)** Expression and spatial localization of *Actin* (*Act5C*) and *Arp53D* from single-cell RNA-seq were visualized using the online *Drosophila* Virtual Expression eXplorer (DVEX)[52]. A threshold of 0.85 was set for each gene.

### Supplementary Tables

**Supplementary Table S1: *Arp53D* orthologs used in phylogenetic analysis**

| Species | NCBI Accession code or Flybase[70] code |
|---|---|
| *D. melanogaster* | FBgn0011743 |
| *D. simulans* | XM_016168248.1 |
| *D. sechellia* | XM_032716929.1 |
| *D. erecta* | FBgn0112814 |
| *D. yakuba* | FBgn0229606 |
| *D. eugracilis* | XM_017223499.1 |
| *D. takahashii* | XM_017160173.1 |
| *D. ficusphila* | XM_017189170.1 |
| *D. anannassae* | XM_001960587.3_modified[a] |
| *S. lebanonensis* | XM_030513294.1 |
| *D. busckii* | XM_017981926.1 |
| *D. mojavensis* | XM_002006572.3 |
| *D. virilis* | FBgn0208134 |
| *D. grimshawi* | XM_001995276.2_modified[a] |
| *D. willistoni* | FBgn0217915 |
| *D. pseudoobscura* | FBgn0078861 |
| *D. persimilis* | XM_002026570.2 |
| *D. miranda* | XM_033397705.1 |

[a]The NCBI gene models were incomplete and were corrected using the BLAT tool[80] in UCSC's genome browser (http://genome.ucsc.edu).

**Supplementary Table S2: RNA-seq databases analyzed**

| Species | Female | Male | Male carcass | Testis |
|---|---|---|---|---|
| *D. melanogaster* | SRR3123319 | SRR3123321 | SRR2021000 | SRR11341471 |
| *D. simulans* | SRR9025064 | SRR9025061 | SRR330567 | SRR9025060 |
| *D. yakuba* | SRR166821 | SRR6161781 | SRR1693754 | SRR934057 |
| *D. anannassae* | SRR7243228, SRR5639307 | SRR6161785 | SRR2021005 | SRR2021004 |
| *D. pseudoobscura* | DRR055272 | DRR055274 | DRR055274 | DRR055270 |
| *D. willistoni* | SRR5639517, SRR7243438 | SRR6161775 | - | SRR7243415, SRR5639494 |
| *D. virilis* | SRR7243394, SRR5639473 | SRR6161774 | SRR5278991 | SRR5278986 |
| *D. mojavensis* | SRR7243269, SRR5639348 | SRR6161773 | - | SRR5639328, SRR7243249 |
| *D. grimshawi* | SRR7253580, SRR3355287 | SRR7253581 | - | SRR3355234, SRR7253527 |
| *S. lebanonensis* | SRR9691967, SRR9691970 | - | SRR9691966 | SRR9691965 |

**Supplementary Table S3: Primers used in this study**

| Purpose | Primer 1's Sequence | Primer 2's Sequence |
|---|---|---|
| Sequencing *Arp53-KO* locus | ACCTTCCCGAATCAAAATCGA | TTCACGTACACCTTGGAGCC |
| Sequencing WT *Arp53D* locus | AGATACTCCCCGTGCTGTCT | GCAAATCCATTGGATCCGCC |
| Testing presence of *Wolbachia*[79] | TTCGCCAATCTGCAGATTAAA | GTTTTAAACGCTTGACAA |
| Sequencing *SOD2* | CTTCAGATCATCGCTGGGCT | TGAAGAATGTTCTGTGCCCGT |
| RT-PCR of Arp53D | ACCTTCCCGAATCAAAATCGA | GCGGCGTGGTGTGAATTAC |

**Supplementary Table S4: Imaging Reagents**

| Antibody or Chemical | Company | Purpose | Dilution |
|---|---|---|---|
| Anti-GFP (chicken) | Abcam (13970) | Western blot | 1:2000 |
| | | Immunofluorescence | 1:500 |
| Anti-tubulin (rabbit) | Abcam (6046) | Western blot | 1:500 |
| | | Immunofluorescence | 1:200 |
| Anti-α-spectrin | Developmental Studies Hybridoma Bank (AB_528473) | Immunofluorescence | 1:50 |
| Anti-phospho-Histone H3 (Ser10) | Millipore (Upstate Brand) | Immunofluorescence | 1:1000 |
| Anti-calmodulin (rabbit) | Gift from Kathleen Beckingham and Leslie Vosshall | Immunofluorescence | 1:50 |
| Anti-mouse Cy3 or Cy5 | Invitrogen | Immunofluorescence | 1:2000 |
| Anti-rabbit Cy3 or Cy5 | Invitrogen | Immunofluorescence | 1:2000 |
| Anti-chicken 488 | Invitrogen | Immunofluorescence | 1:2000 |
| Anti-chicken 680 | LI-COR | Western blot | 1:2500 |
| Anti-rabbit 800 | LI-COR | Western blot | 1:2500 |
| Phalloidin Cy3 | Thermo Fisher | Immunofluorescence | 1:40 |
| Phalloidin Cy5 | Thermo Fisher | Immunofluorescence | 1:40 |
| Sir-actin | Cytoskeleton, Inc | Live imaging | 10 μM |

**Supplementary Table S5:** ***D. melanogaster*** **transgenics constructed**

| Genetic modification | Chromosomal location | Integrated plasmid backbone | Fly strain injected (BDSC) |
|---|---|---|---|
| CRISPR/Cas9 Arp53D knockout | Chr 2, 53D8, 2R:12661915..12662963 | pHD-attP-DsRed (Addgene 51019)[a] | 55821 |
| sfGFP-Arp53D | Chr 3, 89E11, 3R:17052863 | p[acman][81] | 9744 |
| sfGFP-△N-term - Arp53D | Chr 3, 89E11, 3R:17052863 | attB-DsRed[b] | 9744 |
| Nterm Arp53D-sfGFP | Chr 3, 89E11, 3R:17052863 | attB-DsRed[b] | 9744 |
| sfGFP-Nterm Arp53D-Act5C | Chr 3, 89E11, 3R:17052863 | attB-DsRed[b] | 9744 |

[a]pDsRed-attP is from Melissa Harrison & Kate O'Connor-Giles & Jill Wildonger (Addgene plasmid # 51019; http://n2t.net/addgene:51019; RRID:Addgene_51019)
[b]Vector encoding an attB site and 3xP3-*DsRed* flanked by loxP sites.
